# Vitamin B12 attenuates leukocyte inflammatory signature in COVID-19 via methyl-dependent changes in epigenetic marks

**DOI:** 10.1101/2022.08.08.503231

**Authors:** Larissa M. G. Cassiano, Vanessa C. Silva, Marina S. Oliveira, Bárbara V. O. Prado, Cristianne G. Cardoso, Anna C. M. Salim, Gloria R. Franco, Vânia D’Almeida, Saionara C. Francisco, Roney S. Coimbra

## Abstract

COVID-19 induces chromatin remodeling in host immune cells, and it had previously been shown that vitamin B12 downregulates some inflammatory genes via methyl-dependent epigenetic mechanisms. In this work, whole blood cultures from moderate or severe COVID-19 patients were used to assess the potential of B12 as adjuvant drug. The vitamin normalized the expression of a panel of inflammatory genes still dysregulated in the leukocytes despite glucocorticoid therapy during hospitalization. B12 also increased the flux of the sulfur amino acid pathway, raising the bioavailability of methyl. Accordingly, B12-induced downregulation of *CCL3* strongly and negatively correlated with the hypermethylation of CpGs in its regulatory regions. Transcriptome analysis revealed that B12 attenuates the effects of COVID-19 on most inflammation-related pathways affected by the disease. As far as we are aware, this is the first study to demonstrate that pharmacological modulation of epigenetic marks in leukocytes favorably regulates central components of COVID-19 physiopathology.

**Teaser:** B12 has great potential as an adjuvant drug for alleviating inflammation in COVID-19.

## Introduction

Coronavirus disease 2019 (COVID-19), caused by SARS-CoV-2, has been reported, as of August 2022, in more than 583 million confirmed cases and has led to more than 6.4 million deaths worldwide (*1*), although recent studies estimate a much higher death toll (*2*). Since the end of 2020, large-scale population vaccination has drastically reduced the mortality rate and the burden on health systems, but factors such as inequality in access to immunizations, rapid waning neutralizing antibody titers induced either by vaccines or by exposure to SARS -CoV-2 and the emergence of new genotypic variants of the virus have contributed to successive new waves of the pandemic with more dramatic impacts on unvaccinated individuals or those with poor response to vaccines. In this context, efforts have been made in the search for treatments for COVID-19, such as the repositioning of chemicals and monoclonal antibodies that act as anti-inflammatory agents, and the development of new antiviral small molecules that target SARS-CoV-2 replication and antibodies that bind to the viral spike protein blocking virus entry into cells (*3*). Unfortunately, most of these drugs have limited effectiveness or are very expensive, making them unsuitable to be used on a global scale. Therefore, there is still an urgent need for new efficient treatments with safe, inexpensive, and widely available drugs (*4*).

While most COVID-19 patients will experience mild symptoms, some will develop severe acute respiratory distress syndrome, systemic inflammation, multiple organ failure, and other more serious complications that can lead to death. Several components of the host immune system are dramatically altered during SARS-CoV-2 infection and the extent of immune dysregulation is related to the worsening of COVID-19 and progression to death. (*5*). Patients with COVID-19 exhibit inflammatory signatures defined by low levels of type I and III interferons (IFNs) and elevated levels of some cytokines and chemokines. These two branches of the innate immune response contribute to the clinical evolution of the patients (*6*). As the disease worsens, increased levels of inflammatory cytokines and chemokines are observed, which can lead to depletion and exhaustion of T cell populations resulting in a significant elevation of the neutrophil/lymphocyte ratio (NLR) (*7-9*). The increase in neutrophil population (*7, 8*) with transcriptional signatures related to its activated state (*10*) also contributes to the elevation of NLR in COVID-19.

DNA methylation is an epigenetic mechanism that, in mammals, occurs mainly in dinucleotides of a cytosine followed by a guanine (CpG) and can affect the accessibility of transcription factors (TF) and RNA polymerases to DNA, thus modulating gene expression. (*11*). In general, DNA hypomethylation in gene promoters is associated with the activated state of their expression (*12*). It is already known that promoters of genes encoding cytokines are rapidly demethylated (∼6 hours) after T cell activation (*13*) and that several genes encoding key cytokines and chemokines have increased systemic expression in patients with COVID-19 (*14*) are regulated by the methylation of CpGs in their promoters or enhancers (*15-18*). The ability of RNA viruses to hijack the epigenome of host immune cells in order to evade antiviral defense is widely acknowledged (*19, 20*). In early 2021, Corley and colleagues reported that the DNA methylation signature of peripheral blood mononuclear cells from critically ill COVID-19 patients is characterized by hypermethylation of genes related to the IFN-mediated antiviral response and hypomethylation of inflammatory genes (*17*). This finding, in light of previous knowledge about the epigenetic regulation of genes involved in the inflammatory storm of COVID-19, points to the therapeutic potential of drugs capable of modulating the epigenetic landscape as a strategy for the control of exacerbated inflammation resulting from SARS-CoV-2 infection.

The bioavailability of methyl is a determinant of the DNA methylation state and can be regulated by vitamin B12, a cofactor of the enzyme methionine synthase (MS), which transfers methyl groups from 5-methyltetrahydrofolate to homocysteine (HCY) forming methionine. This is then converted to S-adenosylmethionine (SAM), the universal methyl donor. In this same pathway of sulfur amino acids, HCY can be metabolized producing glutathione (GSH) (Fig. 1) (*21*). It has already been demonstrated that adjuvant therapy with B12 increases DNA methylation and reduces the expression of inflammatory mediators in the central nervous system of infant rats with pneumococcal meningitis (*18*). Among these genes, *IL1B* and *CCL3* also play a central role in the pathophysiology of COVID-19. Vitamin B12, also known as cobalamin (Cbl), is an essential micronutrient, which, once absorbed, binds to transcobalamin II (TC-II) an is transported into the cells through specific receptors (*23*). Although the liver is the main source of TC-II, its unsaturated form (not bound to cobalamin) is abundant in the blood (*24*).

**Fig. 1.**
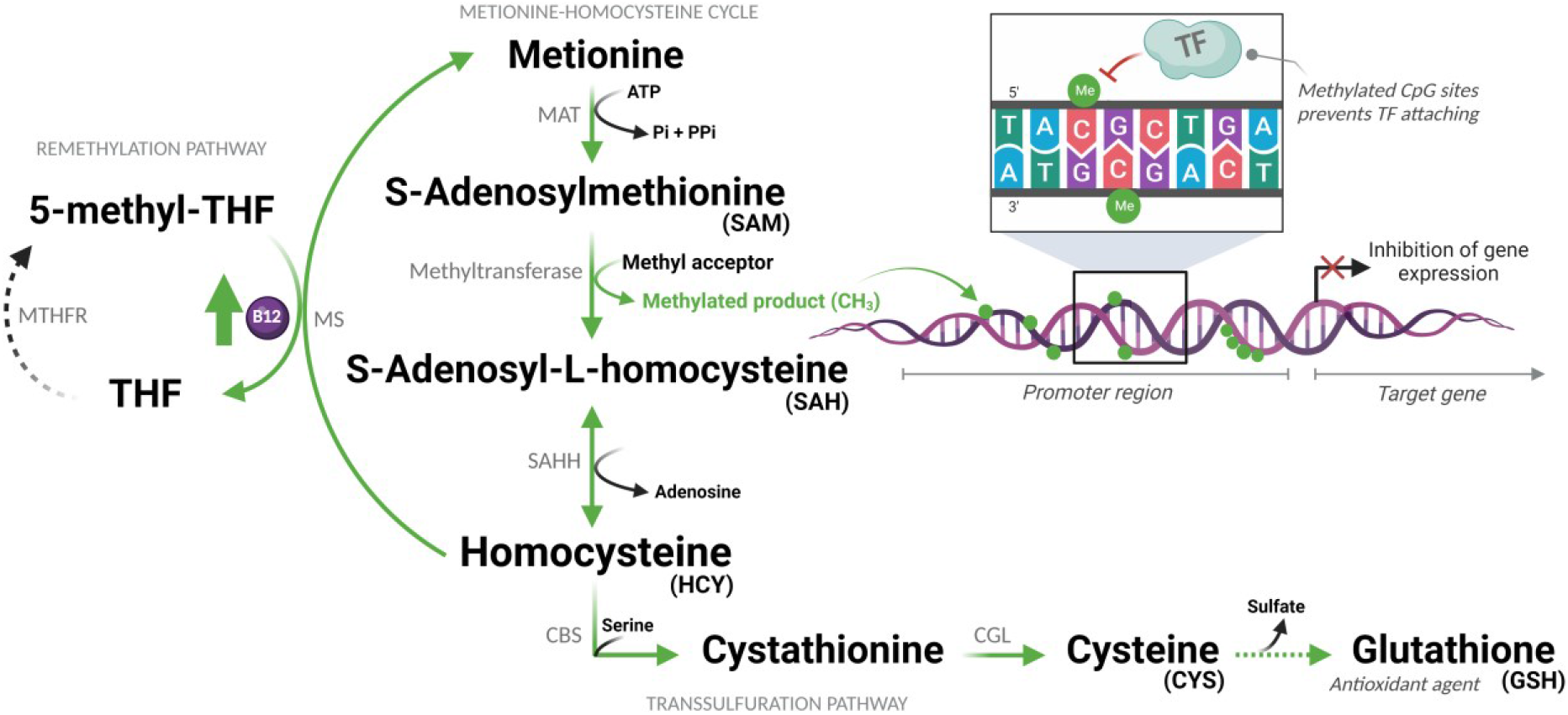
Sulfur amino acid pathway. Methionine is converted to S-adenosylmethionine (SAM), which is the methyl donor for numerous reactions. Upon losing its methyl group, SAM becomes S-adenosyl-L-homocysteine (SAH), which is then converted to homocysteine (HCY). This is then converted back to methionine or enters the transsulfuration pathway to form other sulfur-containing amino acids. Abbreviations: MAT = Methionine adenosyltransferase; ATP = Adenosine triphosphate; Pi = Phosphate (inorganic); PPi = Pyrophosphate; THF = Tetrahydrofolate; MTHFR = Methylenetetrahydrofolate reductase; MS = Methionine synthase; B12 = Vitamin B12; CH_3_/Me = Methyl group; TF = Transcription factors; SAHH = Adenosylhomocysteinase; CBS = Cystathionine beta-synthase; CGL = Cystathionine gamma-lyase. (Created with BioRender.com)

The hypothesis of this study is that, during the advanced phase of COVID-19, characterized by hyperinflammation (cytokine storm), supplemental vitamin B12 would increase the flow of the sulfur amino acid pathway, favoring the production of SAM and the antioxidant GSH. The increased methylation capacity of cells, provided by higher concentrations of SAM, would lead to the hypermethylation of regulatory regions of pro-inflammatory genes attenuating inflammation. This hypothesis was tested using the *ex vivo* model of whole blood culture collected from patients with moderate or severe forms of COVID-19 and healthy controls.

## Results

### Patients and healthy volunteers

The median age of patients and non-infected healthy volunteers included in the study was 64 years (minimum = 45; maximum = 86), 55% of which were women, with no statistically significant differences between the groups in these aspects. All patients had a confirmatory clinical diagnosis of COVID-19 and confirmation of SARS-CoV-2 virus infection by RT-qPCR carried out, on average, 5 days before the collection of blood samples for this project. The time elapsed between the onset of symptoms and admission to the hospital was 6.3 days, and the length of stay before sample collection was 11.8 days, for patients in the MOD and SEV groups, with no statistically significant difference between the two groups. Cardiovascular diseases (73.08%) and diabetes mellitus (38.46%) were the most frequent comorbidities among patients.

All patients were on glucocorticoid treatment (dexamethasone: 25 patients; prednisone: 1 patient) but those with severe COVID-19 most often received combinations of two or three drugs of this category, namely dexamethasone, hydrocortisone and beclomethasone for an average of 11 days before their blood samples were taken, with no differences between MOD and SEV regarding the duration of this treatment. Among the cytochemical parameters evaluated, differences were found between MOD and SEV groups for blood glucose (mg/dL; means MOD = 131.7 and SEV = 196.7; *P* = 0.0111), total leukocytes (cells per mm^3^; means MOD = 9,650 and SEV = 16,355; *P* = 0.0041), percentage of lymphocytes (%; medians: MOD = 14.85 and SEV = 6.95; *P* = 0.0048), percentage of neutrophils (%; medians: MOD = 73.85 and SEV = 86.1; *P* = 0.0139) and for the NLR (means MOD = 5.2 and SEV = 13.1; *P* = 0.0038). The increase in the percentage of neutrophils in SEV patients coincides with the occurrence of more frequent bacterial coinfections in this group (20% for MOD and 75% for SEV; *P* = 0.0138). Arterial blood gas analysis of MOD and SEV patients revealed significant differences in pH (medians MOD = 7.435 and SEV = 7.375; *P* = 0.0223), O_2_ pressure (mmHg; medians MOD = 58.15 and SEV = 80.10; *P* = 0.0028), CO_2_ tension (mmol/L; medians MOD = 20.30 and SEV = 23.25; *P* = 0.0309) and O_2_ saturation (%; means MOD = 90.25 and SEV = 94.94; *P* = 0.0123), compatible with the fact that SEV patients were intubated with mechanical ventilation. Potassium concentrations were also higher in patients in the SEV group (mmol/L; medians MOD = 3.820 and SEV = 4.365; *P* = 0.0022). The combination of Clavulanate and Amoxicillin was more frequent among MOD patients than SEV (%; MOD = 70 and SEV = 25; *P* = 0.0426). Although there were no statistically significant differences between MOD and SEV groups regarding the occurrence of clinical complications, the outcomes were significantly worse for patients in the SEV group, with longer total hospitalization time (days; means MOD = 16.2 and SEV = 27.94; *P* = 0.0132) and more frequent deaths (%; MOD = 10% and SEV = 75%; *P* = 0.0036). All information compiled from medical records is presented in Table S1.

Control subjects had no RT-qPCR detectable SARS-CoV2 in their oropharynx and nasopharynx and no antibodies against the virus in their blood. No statistically significant differences were found between baseline plasma B12 levels of patients in the SEV, MOD and CTRL groups (Fig. S1). No SARS-CoV-2 RNA was detected by RT-qPCR in raw blood or blood cultures of any participant in this study.

### Validation of the experimental model

The RT-qPCR analyses revealed distinctive transcriptional signatures for MOD and SEV forms of COVID-19 and the CTRL group in the aliquots of endpoint Z, which remained mostly preserved after 24h of incubation (endpoint A) (Fig.S2 and Fig. 2). Importantly, patients were already on glucocorticoid treatment before blood collection for this study, which may explain why higher levels of mRNA were not found for some genes, such as *CCL2, CXCL9, IL6, IL17A, CCL1* and *TNF* in MOD and/or SEV patients compared to CTRL at endpoints Z and A (Fig.S2 and Fig. 2). High levels of *CCL3* and *IL1B* mRNA were observed in the SEV and MOD groups compared to the CTRL, indicating that these mediators of COVID-19 inflammation are not sufficiently responsive to glucocorticoid treatment. Reduced levels of mRNA of marker genes of CD4 and CD8a lymphocyte lineages were observed in MOD and SEV patients compared to CTRL at endpoints Z and A. At endpoint Z, these two genes showed a positive correlation (MOD: r = 0.5553, *P* = 0.004 and SEV: r = 0.5665, *P* = 0.0032) with the percentage of lymphocytes in the patients’ blood counts (Table S1). On the other hand, mRNA levels of *HAVCR2* lymphocyte exhaustion marker did not differ between patients and CTRL.

**Fig. 2.**
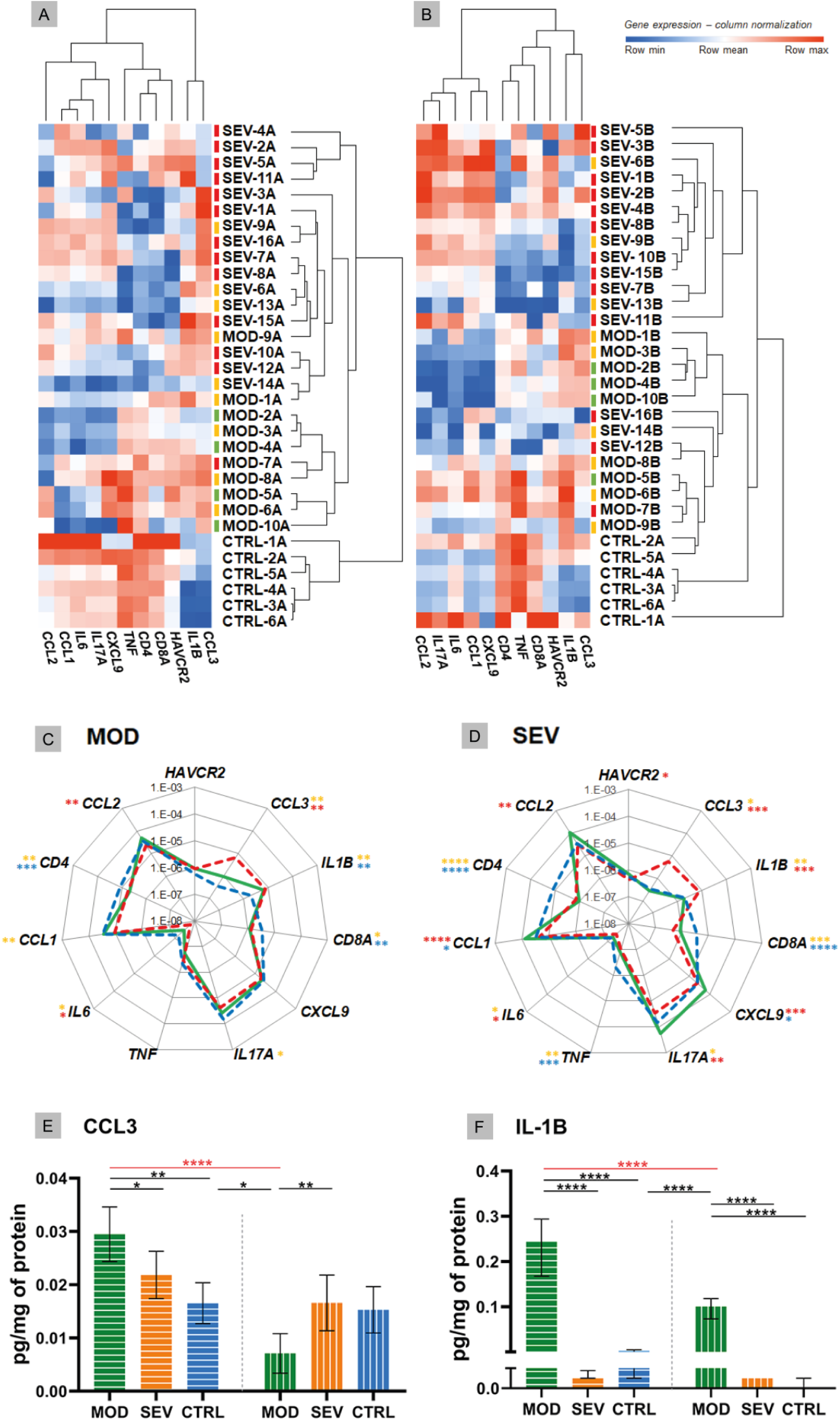
Vitamin B12 favorably modulated critical inflammatory mediators. **A** and **B**: Dendrogram and heatmap depicting the hierarchical clustering and expression levels of COVID-19 hyperinflammation-related genes panel in whole blood cultures of untreated MOD and SEV patients (endpoint A) (Panel A) or treated with B12 (endpoint B) (Panel B) and untreated non-infected controls (endpoint A). Colored dashes next to the sample identification correspond to the patient’s outcome. Green dashes: patients discharged from hospital one day after sample collection; Yellow lines: patients discharged from hospital two or more days after sample collection; Red lines: patients died. **C** and **D**: Radar charts depicting the expression levels of COVID-19 hyperinflammation-related genes panel in whole blood cultures. The filled green lines correspond to gene expression values of infected groups (MOD, Panel C or SEV, Panel D) at endpoint B. Dashed red lines correspond to gene expression values of infected groups at endpoint A. Dashed blue lines correspond to gene expression values control group at endpoint A. The gene expression values (2e(-ΔCt)) of the groups were compared pairwise using two-tailed Student’s t, paired Student’s t, Mann-Whitney and Wilcoxon tests according to experimental design and data distribution. Values were expressed as medians. N sample = MOD (10) and SEV (16) for each endpoint (A or B), CTRL (6). Yellow asterisks = Infected A versus controls A; Red asterisks = Infected B versus Infected A; Blue asterisks = Infected B versus controls A. **E** and **F**: Bar charts depicting intracellular protein levels of CCL3 (Panel E) and IL-1B (Panel F). Filled bars with horizontal lines = endpoint A. Filled bars with vertical lines = endpoint B. Data were compared using two-way Analysis of Variance (ANOVA) test followed by Tukey’s multiple comparison test. Values were represented as mean ± standard deviation. N sample = MOD (7), SEV (8), CTRL (4) for each endpoint (A or B). Red asterisks and red lines denote the effect of B12 on cultures of patients with moderate COVID-19. * *P* < 0.05; ** *P* < 0.01; *** *P* < 0.001; **** *P* < 0.0001. Abbreviations: SEV = severe COVID-19; MOD = moderate COVID-19; CTRL = non-infected controls. Suffixes: A = endpoint A (samples added to culture medium with excipient and incubated for 24h); B = endpoint B (samples added to culture medium with B12 and incubated for 24h).

### Vitamin B12 favorably modulated critical inflammatory mediators

The biomarker genes *CCL3* and *IL1B*, which were upregulated in patients despite previous glucocorticoid therapy, responded very well to treatment with B12, which reduced *CCL3* mRNA levels of MOD and SEV, and *IL1B* of SEV, matching them to the CTRL at endpoint A (Fig. 2, A to D). Importantly, B12 did not affect *CCL3* or *IL1B* expression in the CTRL group at endpoint B. Other genes had lower mRNA levels in MOD (*IL17A, IL6* and *CCL1*) and SEV (*IL17A* and *IL6*) compared to CTRL at endpoint A. B12 raised mRNA levels of *IL6* and *IL17A* in both MOD and SEV, matching them to CTRL at endpoint A. The vitamin also raised *CCL1* mRNA levels in cultures of MOD and SEV patients. In this case, *CCL1* mRNA levels of MOD group equaled those of the CTRL at endpoint A and the levels of the SEV exceeded this reference. *CXCL9* mRNA levels, which did not differ between patients and CTRL at endpoint A, increased in SEV cultures treated with B12 so that its mRNA levels exceeded those of the CTRLs at endpoint A.

*TNF*, whose baseline mRNA levels were low only in the SEV group compared to CTRL at endpoint A, did not have their mRNA levels significantly altered by B12 in either of the two patient groups. B12 also did not affect *CD4* and *CD8A* mRNA levels in MOD or SEV, but slightly increased those of *HAVCR2* in SEV. However, this increase was not enough to achieve a statistically significant difference compared to CTRL at endpoint A.

Finally, the treatment of whole blood cultures of individuals in the CTRL group with B12 did not change the expression levels of any gene assessed. Overall, these results reinforce the security of the 1 nM dose of B12 and prove its efficiency in regulating the mRNA levels of several critical inflammatory mediators in whole blood cultures of patients with moderate and severe COVID-19. This regulation of mRNA levels of inflammatory mediators by B12 was done by down- or upregulation depending on the gene.

### Vitamin B12 reduced intracellular protein levels of critical inflammatory mediators

MOD cultures treated with B12 had reduced intracellular protein levels of CCL3 and IL-1B (*P* < 0.0001), both of which were elevated at endpoint A compared to CTRL (*P* = 0.0011 and *P* < 0.0001, respectively) (Fig. 2, E and F). Interestingly, B12 did not affect the intracellular protein levels of IL-1B in the cultures of the SEV group, and, regardless of the treatment, the intracellular protein levels of IL-1B were very low, close to or below the detection limit of the method (Fig. 2F).

### Vitamin B12 increased the flow of the sulfur amino acid pathway

B12 caused an increase in HCY concentrations of all groups (MOD, *P* = 0.0037; SEV, *P* < 0.0001; CTRL *P* = 0.0313) and in CYS and GSH levels of MOD and SEV (MOD: *P* = 0.0029 and *P* = 0.0002; SEV: *P* = 0.0007 and *P* < 0.0001) compared to their respective levels at endpoint A, which indicates an increased flow of the sulfur amino acid pathway (Table 1). However, the expected increase in the SAM/SAH ratio in response to B12 was not observed. In fact, at endpoint B, decreased SAM levels were observed in MOD group (*P* = 0.0010), without, however, any change in SAM/SAH ratio. In the SEV group, B12 increased SAH (*P* = 0.0091) and reduced SAM/SAH (*P* = 0.0021).

**Table 1.** Vitamin B12 increases the flow of the sulfur amino acid pathway.

| Metabolites | Group | Intracellular concentrations (μmol/g Hb)<br>(mean [SD] or median [IQ]) |  |  | P value (AxB) |  |  |
| --- | --- | --- | --- | --- | --- | --- | --- |
|  |  | MOD | SEV | CTRL | MOD | SEV | CTRL |
| HCY | A | 0.04096<br>(0.007656) | 0.04312<br>(0.01524) | 0.054<br>(0.05133-0.0638) | 0.0037<br>(**) | <0.0001<br>(****) | 0.0313<br>(*) |
|  | B | 0.0561<br>(0.01653) | 0.05762<br>(0.01784) | 0.067<br>(0.06225-0.0985) |  |  |  |
| CYS | A | 0.5869<br>(0.2176) | 0.6795<br>(0.2867) | 0.5635<br>(0.04167) | 0.0029<br>(**) | 0.0007<br>(***) | 0.1253 |
|  | B | 0.7326<br>(0.2542) | 0.9097<br>(0.3812) | 0.8457<br>(0.2764) |  |  |  |
| GSH | A | 12.52 (2.123) | 10.6 (4.292) | 15.35 (2.735) | 0.0002<br>(***) | <0.0001<br>(****) | 0.0612 |
|  | B | 15.1 (2.877) | 14.94 (6.213) | 16.24 (2.878) |  |  |  |
| SAM | A | 0.002436<br>(0.000673) | 0.003405<br>(0.00225-0.00466) | 0.0018<br>(0.001418-0.001815) | 0.0010<br>(**) | 0.4954 | 0.0938 |
|  | B | 0.001789<br>(0.0003617) | 0.002673<br>(0.002365-0.004239) | 0.002081<br>(0.001825-0.00227) |  |  |  |
| SAH | A | 0.0002819<br>(0.00009389) | 0.0001972<br>(0.00008667) | 0.0001817<br>(0.00003608) | 0.0722 | 0.0015<br>(**) | 0.0087<br>(**) |
|  | B | 0.0002293<br>(0.00004081) | 0.0003267<br>(0.0001541) | 0.0002277<br>(0.00004052) |  |  |  |
| SAM:SAH | A | 9.093 (2.571) | 16.44 (12.09) | 9.632 (3.123) | 0.1511 | 0.0021<br>(**) | 0.7341 |
|  | B | 8.041 (2.28) | 8.058 (4.557) | 9.243 (2.087) |  |  |  |

The parametric data were compared using two-tailed paired Student's T test and represented as ± standard deviation. The nonparametric data were compared using two-tailed Wilcoxon paired test and represented as ± interquartile range. \* *P* < 0.05; \*\* *P* < 0.01; \*\*\* *P* < 0.001 \*\*\*\*; *P* < 0.0001. Abbreviations: SEV = severe COVID-19; MOD = moderate COVID-19; CTRL = non-infected controls; A = endpoint A (samples added to culture medium with excipient and incubated for 24h); B = endpoint B (samples added to culture medium with B12 and incubated for 24h).

### Vitamin B12 increased methylation levels of CpGs in regulatory regions of *CCL3*

The effects of COVID-19 and B12 on the methylation levels of the 21 CpGs located in the promoter region and proximal portion of the first exon of the *CCL3* gene (GRCh38/hg38 chr17: 36,090,276-36,090,005) were evaluated by BSP in an NGS platform from DNA libraries produced with aliquots of endpoints A and B from individuals representing MOD, SEV and CTRL groups. At endpoint A, when compared with the CTRL group, no changes were found in the percentage of methylation of any of the evaluated positions in the cultures of MOD patients, while SEV patients had an hypomethylated CpG at chr17:36,090,102 (Fig. 3). Treatment of cultures with B12 increased methylation levels of CpGs at chr17:36,090,097 in MOD subjects and at chr17:36,090,097, 36,090,102 and 36,090,246 in SEV subjects compared to their respective cultures at endpoint A. Note that for methylation of the chr17:36,090,102 position in SEV group, COVID-19 and B12 had inverse effects. The methylation levels of all the aforementioned CpGs had negative and statistically significant correlations with the gene expression levels.

**Fig. 3.**
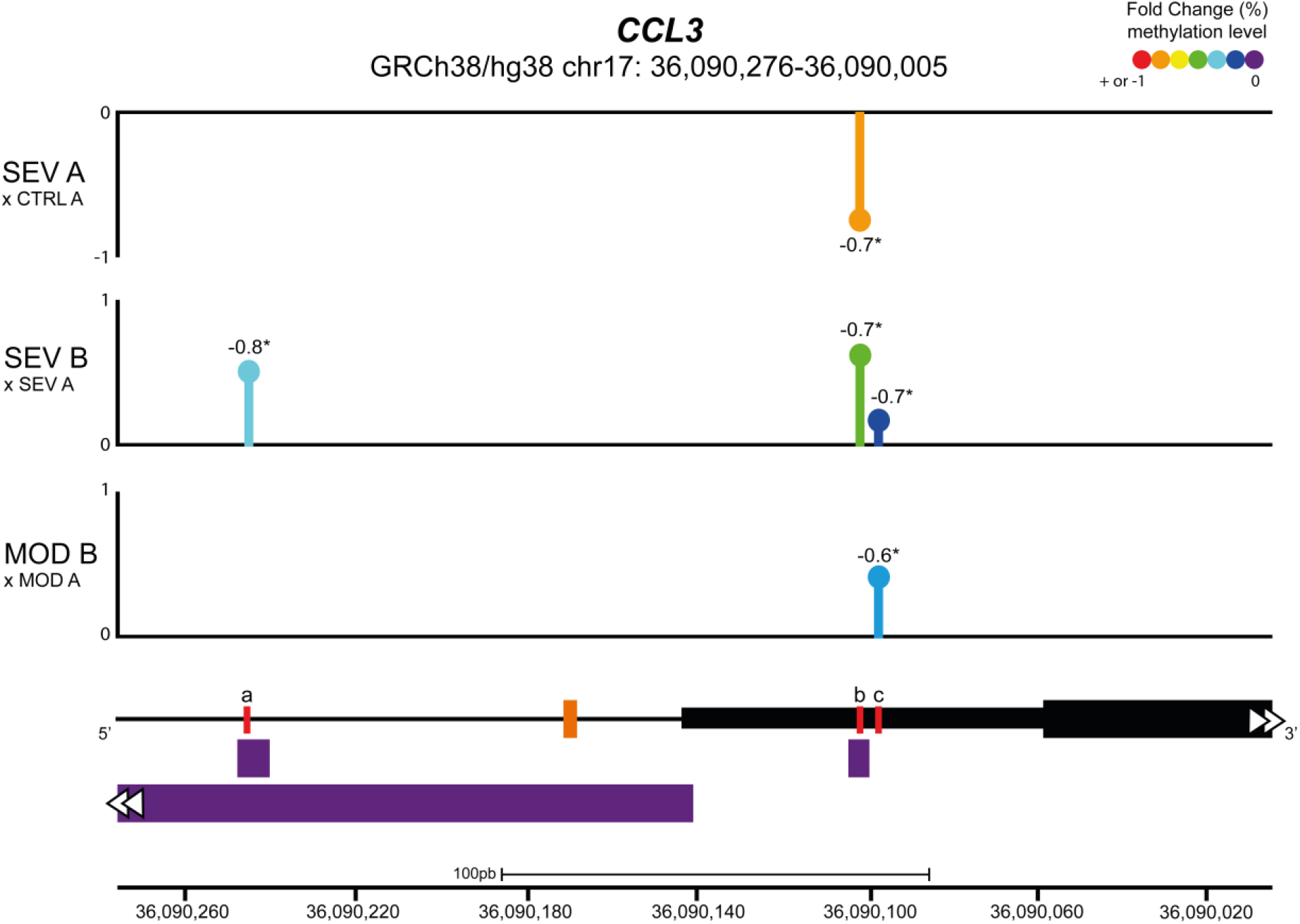
Vitamin B12 increased methylation levels of CpGs in regulatory regions of *CCL3*. Lollipop plot depicting the percent fold change of methylation levels of CpG sites in the promoter region and proximal portion of the first exon of the *CCL3* gene. Values next to lollipops represent correlation coefficients (Pearson or Spearman) with statistical significance (* *P* < 0.05) between methylation levels in each CpG position and gene expression values. The % fold change of methylation levels of CpG positions were compared pairwise using two-tailed Mann-Whitney. N sample = MOD (5) and SEV (5) for each endpoint (A or B), CTRL (5). In the schematic representation of the analyzed gene region, the black line represents the *CCL3* promoter region and the black blocks delimit the 5’ UTR region (thin block) and proximal portion of the first exon of *CCL3* (thick block). The orange dash indicates the transcription start site (TSS) of *CCL3* and the purple blocks indicate transcription factor binding sites (TFBS). Red lines correspond to differentially methylated CpG sites, coordinates (GRCh38/hg38): a = chr17:36,090,246; b = chr17:36,090,102; c = chr17:36,090,097. Abbreviations: SEV = severe COVID-19; MOD = moderate COVID-19; CTRL = untreated non-infected controls. Suffixes: A = endpoint A (samples added to culture medium with excipient and incubated for 24h); B = endpoint B (samples added to culture medium with B12 and incubated for 24h).

### Vitamin B12 attenuated the pro-inflammatory profile of leukocytes from patients with COVID-19

The effects of COVID-19 and B12 on the global gene expression in leukocytes in whole blood cultures from representative individuals of the MOD, SEV and CTRL groups were assessed by RNA-Seq. Aiming at identifying differentially expressed genes (DEG), the following contrasts were analyzed: 1) MOD vs. CTRL at endpoint A; 2) SEV vs. CTRL at endpoint A; 3) MOD (endpoint B) vs. CTRL (endpoint A); and 4) SEV (endpoint B) vs. CTRL (endpoint A); 5) CTRL at endpoint A vs. CTRL at endpoint B.

A large number of genes had their expression affected by COVID-19 (Fig. 4). In the MOD group, 3,034 DEGs (2,041 upregulated and 993 downregulated) were found in contrast 1 and 3,636 DEGs (2,361 upregulated and 1,275 downregulated) in contrast 3. In cultures of the SEV group, 8,565 DEGs (4,464 upregulated, 4,101 downregulated) were found in contrast 2 and 8,894 DEGs (4,520 upregulated and 4,374 downregulated) in contrast 4. Among the DEGs identified, 2,699 had decreased (1,364) or increased (1,335) expression after treatment with B12 (Fig. 4). In the leukocytes of patients in MOD group, B12-upregulated genes had GO annotations related to *phagocytosis, regulation of miRNA transcription, response to virus* and *monocyte extravasation*. In the same group, B12-downregulated genes had GO annotations related to *NF-kappaB signaling, T-cell receptor signaling pathway* and *negative regulation of histone H3-K9 trimethylation* (Fig. S3). Regarding the SEV group, B12-upregulated genes had GO annotations related to *cytoplasmic translation* (tRNA and rRNA processing), *adaptive immune response, ncRNA processing, miRNA-mediated gene silencing, proteasome-mediated ubiquitin-dependent protein catabolic process, cobalamin transport*, and *negative regulation of transcription by RNA polymerase II* (Fig. S3). B12-downregulated genes in this group were related to *positive regulation of transcription by RNA polymerase II, processing* and *transport of mRNA, mRNA splicing, cellular response to DNA damage stimulus, activation of innate immune response, positive regulation of IL-6* and *IL-2 production, T-cell differentiation in thymus, histone modification, viral transcription* and *DNA conformational change*. Finally, no DEGs were found in contrast 5, which reinforces the safety profile of B12.

**Fig. 4.**
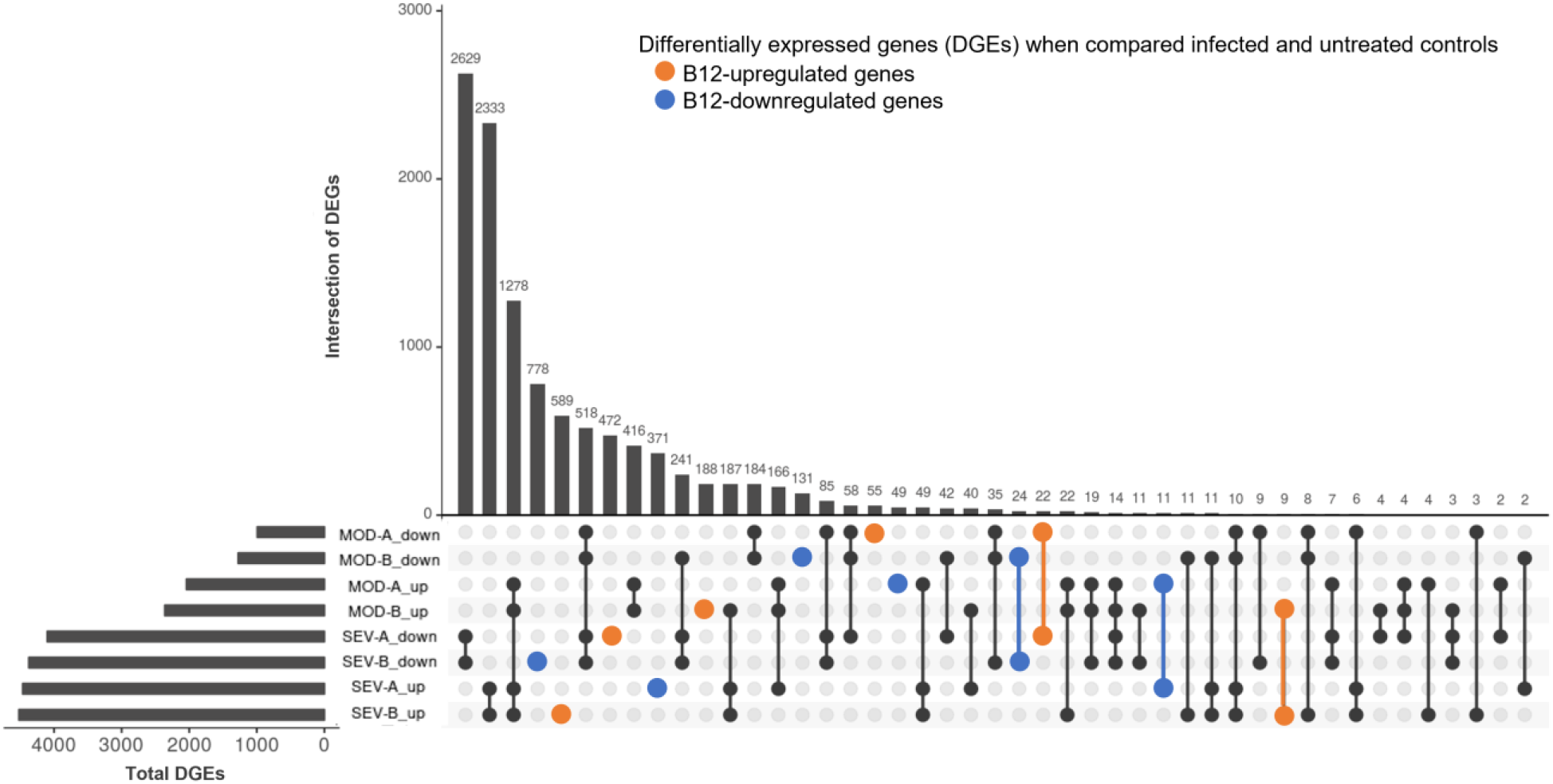
Differently expressed genes (DEGs). DEGs in infected patients (at endpoint A or B) when compared to untreated non-infected controls (endpoint A). Abbreviations: SEV = severe COVID-19; MOD = moderate COVID-19. Suffixes: A = endpoint A (samples added to culture medium with excipient and incubated for 24h); B = endpoint B (samples added to culture medium with B12 and incubated for 24h); up = upregulated; down = downregulated.

Functional enrichment analysis of DEGs identified in the contrasts above mentioned revealed 90 metabolic and signaling pathways differentially regulated in the groups MOD or SEV at endpoint A directly or indirectly related to the inflammatory response in blood, although patients had received glucocorticoid treatment for approximately 11 days prior to collection of samples to the study (Fig. 5A and Fig. S4). In the cultures treated with B12, 45 and 74 out of these 90 pathways were still differentially regulated in MOD and SEV groups, respectively, although with a minimum 20% difference in Z-scores relatively to their untreated condition. An overall favorable effect on inflammation control was predicted for B12 treated cultures of MOD (Fig. 5B and Fig. S5) and SEV (Fig. 5C and Fig. S6) groups.

**Fig. 5.**
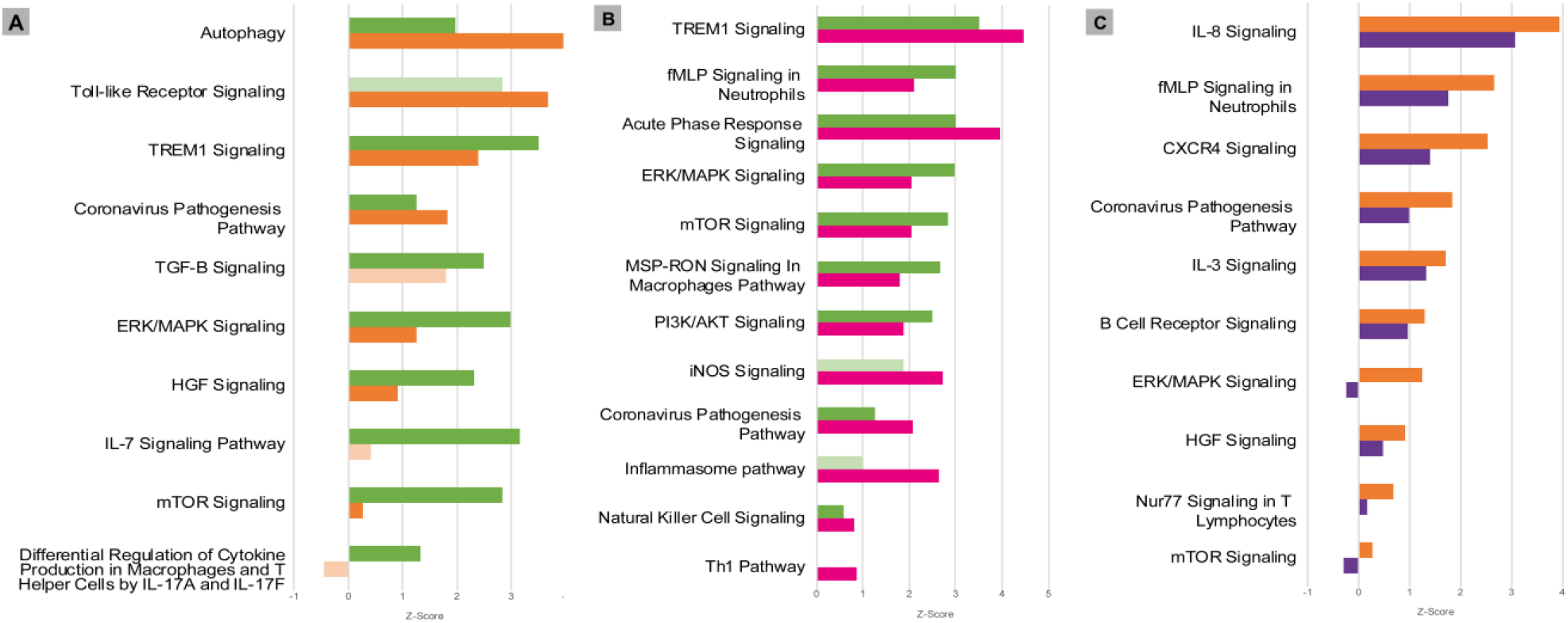
Canonical Pathways affected by COVID-19 and B12. Patients with moderate and severe COVID-19 previously treated with glucocorticoids had distinct global gene expression patterns **(A)** and vitamin B12 attenuated the pro-inflammatory profile of leukocytes from patients with moderate **(B)** or severe **(C)** forms of the disease. Green bars = contrast 1 (MOD vs. CTRL at endpoint A); Orange bars = contrast 2 (SEV vs. CTRL at endpoint A); Pink bars = contrast 3 (MOD at endpoint B vs. CTRL at endpoint A); Purple bars = contrast 4 (SEV at endpoint B vs. CTRL at endpoint A); Faded bars indicate pathways with *P* value less than 0.05 (statistically non-significant). Abbreviations; SEV = severe COVID-19; MOD = moderate COVID-19. Suffixes: A = endpoint A (Samples added to culture medium with excipient and incubated for 24h); B = endpoint B (Samples added to culture medium with B12 and incubated for 24h).

## Discussion

Epigenetic changes in host cells during COVID-19 have already been described and are associated with the regulation of the SARS-CoV-2 cycle (*25, 26*), the disease severity (*17, 26*) or the prognosis for critically ill patients (*16, 26*). However, to the best of our knowledge, this is the first work to demonstrate that pharmacological modulation of the leukocyte epigenetic marks favorably regulates central components of hyperinflammation in moderate and severe forms of the disease. Initially, the whole blood culture model was validated for evaluation of anti-inflammatory drugs for COVID-19 by the identification of stable transcriptional signatures distinguishing between moderate and severe forms of the disease and between these and controls without infection, despite the fact that patients were on glucocorticoids before sample collection. Then, using this *ex vivo* model, it was demonstrated that vitamin B12 favorably modulates, by methyl-dependent epigenetic mechanisms, the expression of inflammatory genes and the activity of metabolic and signaling pathways related to the hyperinflammation associated with COVID-19 forms that require hospitalization.

Expression analysis of a panel of COVID-19 related genes at endpoint Z (blood aliquots added with culture medium and B12 excipient but not incubated) (Fig. S2) in addition to revealing distinctive transcriptional signatures of moderate and severe forms of the disease, have also shed light on the effects of glucocorticoid therapy on circulating leukocytes. Exacerbated activation of inflammatory cytokines is associated with COVID-19 severity and poor prognosis (*27*). However, at endpoint Z, mRNA levels of *CCL2, CXCL9, IL6, IL17A, CCL1* and *TNF* were lower in the MOD and/or SEV groups compared to the controls, probably due to previous glucocorticoid treatment that patients received during hospitalization. Although these genes are components of the hyperinflammation caused by SARS-CoV-2 (*27, 28*), it is debatable whether their early inhibition and drastic reduction are beneficial for patients. For instance, dexamethasone is known to have a strong inhibitory effect on *IL6* and *IL17A* in patients with COVID-19 (*28*), but it is still unclear to what extent the beneficial effects of IL-6 blockers depend on dose, time of administration, clinical condition, among other factors (*29, 30*). It is also worth mentioning that IL-17A play a role in induce protective inflammatory responses, hindering viral infection (*31, 32*). In contrast, some biomarkers remained overexpressed in MOD and SEV cultures, despite the patients having been previously treated with glucocorticoids. Among the genes most refractory to glucocorticoid treatment, *CCL3* and *IL1B* stand out, both with a central role in the pathophysiology of COVID-19 and highly expressed in peripheral blood mononuclear cells (PBMC) during the disease (*14, 33*). CCL3 is a critical pyrogenic cytokine that is involved in leukocyte recruitment and activation in acute inflammation. It has been reported that *CCL3* expression is higher in COVID-19 patients with an unfavorable outcome (*34*). IL-1B has a broad spectrum of biological functions and participates in innate and adaptive immunity. In infections, IL-1B induces gene expression and synthesis of various cytokines and chemokines in macrophages and mast cells. SARS-CoV-2 activated IL-1B stimulates the secretion of TNF, IL-6 and other cytokines, a pro-inflammatory complex that can lead to cytokine storm and be deleterious both in the lung and systemically (*35*).

The transcriptional signatures observed at endpoint Z remained for the most part conserved after incubation of cultures for 24h (endpoint A), thus validating the *ex vivo* model of whole blood cultures for researching drugs with the potential to fight hyperinflammation in COVID-19.

The whole blood culture model was used to test whether vitamin B12 can regulate, via methyl-dependent epigenetic mechanisms, the expression of inflammatory genes in the leukocytes of patients with moderate and severe forms of the disease and who are already being treated with glucocorticoids. Indeed, in the cultures of the MOD group, B12 equaled the mRNA levels of almost all the cytokine or chemokine genes tested to those of the CTRL group that did not receive the vitamin. These genes had not been regularized by glucocorticoid therapy previously received by the patients before sample collection to this study. An exception was *IL1B*, whose mRNA levels were not significantly affected by B12. Interestingly, B12 decreased intracellular protein levels of IL-1B in the MOD group (Fig. 2F). It is worth noting the magnitude of reduction of *CCL3* mRNA levels by B12 in this group, which was accompanied by the reduction of their intracellular protein levels. Likewise, the treatment of SEV group cultures with B12 brought the mRNA levels of most of the evaluated biomarkers closer to those of untreated CTRLs (Fig. 2, A, B and D). In this group, *CCL3* and *IL1B* were the most responsive genes, at the transcriptional level, to B12, but no differences were observed in the intracellular levels of the respective proteins (Fig. 2, E and F). The choice to analyze the intracellular protein concentrations of IL-1B and CCL3 was based on the premise that the inhibition of gene transcription in response to B12 added to the turnover of these proteins by the proteasome machinery present inside the cells would result in detectable reductions in their concentrations after 24 hours of culture incubation. As opposite, in the extracellular medium, a detectable decrease in the concentrations of these proteins would be less likely due to the absence of the cellular machinery for protein degradation. This strategy was suitable to detect the reduction of CCL3 and IL-1B proteins in cultures of the MOD group. However, in cultures of the SEV group, the reduction of CCL3 by B12 did not reach statistical significance and intracellular IL-1B levels were very low, close to the detection limit of the technique, suggesting that, in severe COVID-19, leukocytes are more prone to promptly export IL-1B. Still in the SEV group, B12 increased the mRNA levels of *CCL1* and *CXCL9* beyond those observed in the untreated CTRL group. CCL1 is produced by activated T lymphocytes and monocytes/macrophages, and is the only known chemokine capable of interacting with CCR8, a receptor expressed on Th2 cells and regulatory T cells (Treg). This chemokine mediates the inhibition of dexamethasone-induced thymocyte apoptosis (*36*) and its stimulation is related to autocrine anti-apoptosis (*37*), suggesting that CCL1-CCR8 interactions may provide survival signals for T cells at sites of inflammation. CXCL9, which is inducible by IFN-γ, is related to the activation of the Th1 antiviral immune response, as well as the trafficking of Th1, CD8 and natural killer (NK) cells (*38*). B12 treatment also brought *IL17A* levels of SEV closer to those of untreated CTRL. Therefore, these results indicate that B12 can stimulate T cell survival in patients with severe COVID-19. As expected, since lymphocytes expressing *CD4* and *CD8A* multiply and differentiate by clonal expansion in the thymus (*39*), treatment of whole blood cultures with B12 did not affect *CD4* and *CD8A* mRNA levels in either MOD or SEV groups (Fig. 2, A to D).

Quantification of sulfur amino acid pathway metabolites revealed an increased flux as evidenced by higher concentrations of HCY, CYS and GSH in the cultures of the MOD and SEV groups treated with B12 compared to their untreated cultures. The increased concentration of GSH favors the attenuation of exacerbated inflammation and prevents cell damage caused by COVID-19 due to its antioxidant effect (*40*). However, a reduction in SAM and SAM/SAH was also observed in cultures from MOD and SEV groups, respectively, in response to B12 (Table 1). When HCY and B12 levels are high, the enzyme MS converts HCY to methionine, which is then converted to SAM. Then, methyltransferases transfer the methyl group of SAM to cytosine residues in the DNA or to other acceptors, such as amino acid residues in histone tails, resulting in the formation of SAH (*41*). In fact, an increase in SAH was observed in cultures of the SEV group treated with vitamin B12. This set of results suggests that SAM, whose concentration was probably increased in cultures treated with B12, was readily consumed in methylation reactions of either DNA or other acceptors of methyl groups during the incubation period of 24 h. This increase in the methylation capacity of leukocytes induced by B12 was confirmed by analyzing the methylation profile of CpGs of the *CCL3* gene.

Leukocytes from cultures of patients with severe COVID-19 at endpoint A showed hypomethylation of the chr17:36,090,102 locus, in the 5’ UTR region of *CCL3*, when compared to the CTRL group (Fig. 3). Contrarily, no DML was identified in leukocyte cultures from patients in the MOD group at endpoint A. The hypothesis that B12 downregulates inflammatory genes of COVID-19 via methyl-dependent epigenetic mechanisms was proved by the hypermethylation of three CpGs located in the 5’ UTR region and in the proximal portion of the first exon of *CCL3* (Chr17: 36,090,276 - 36,090,005) of leukocytes from cultures of MOD and SEV groups treated with the vitamin. The B12-induced increment in methylation levels of each of these DMLs negatively correlated with the *CCL3* expression level assessed by RT-qPCR (Fig. 3). Two of the three DMLs in response to B12 are located at transcription factor binding sites (TFBS). TFs FOS (JASPAR ID: MA1800.1) and SMARCA4 (ORegAnno ID: OREG1238466) bind to the region encompassing the DML chr17:36,090,246 (a), while NFIC (JASPAR ID: MA0161.2) binds to that containing the DML chr17:36,090,102 (b). NFIC is a component of the CTF/NF-I family and a SARS-CoV2-upregulated inflammation-associated TF (*42*). At endpoint A, the gene encoding NFIC is upregulated in the SEV group compared to controls (contrast 2), suggesting an important role for this TF in the transcription of inflammatory genes in severe COVID-19. AP-1, a complex formed by FOS and other TFs, is crucial for transcription of *CCL3*. The AP-1 signaling pathway can be activated by cytokines, growth factors, stress, and bacterial or viral infections, including SARS-CoV proteins (*43*). Interestingly, the gene encoding FOS was downregulated in the SEV group at endpoint B compared to controls at endpoint A (contrast 4), indicating that B12, in addition to blocking the binding of this TF to the regulatory region of *CCL3*, also reduces its expression in severe COVID-19. SMARCA4 is a protein present in ATP-dependent chromatin remodeling complexes of the SWI/SNF type. Members of this family have helicase and ATPase activities and regulate the transcription of certain genes by altering the chromatin structure in their surroundings. Recruitment of this complex is associated with increased expression of *CCL3* (*44*).

Therefore, the severe form of COVID-19 induces hypomethylation of a CpG locus in the regulatory region of *CCL3*, which is coherent with its upregulation. This result corroborates the findings of Corley et al (*17*), who had already demonstrated the hypomethylation of upregulated inflammatory genes in PBMC of patients with severe COVID-19. More importantly, the results presented herein prove that vitamin B12 downregulates *CCL3* in leukocytes of patients with moderate or severe COVID-19 via the hypermethylation of CpGs and subsequent inhibition of TF binding to the analyzed regulatory region.

As expected, functional enrichment analysis of RNA-Seq data revealed different immune response profiles depending on the severity of the disease (Fig. 5A and Fig. S4). However, although it has already been shown that greater activation of hyperinflammation-related pathways is associated with disease severity (*45*), in the present study, cultures from patients with severe COVID-19 showed less activation or greater inhibition of pathways directly or indirectly related to inflammation compared to patients with moderate disease. It is very likely that the less inflamed profile of SEV group is due to the administration of more potent glucocorticoids to these patients during hospitalization, prior to the collection of blood samples for the study (Table S1). These differences in the inflammatory profile of leukocytes of patients with severe and moderate COVID-19 previously treated with glucocorticoids are evident in the activation patterns of the *Coronavirus Pathogenesis Pathway* (Fig. S7).

Gene ontology (GO) functional annotation analysis of the 2,699 genes differentially expressed in response to B12 (Fig. S3) indicate that, in leukocytes of patients with moderate COVID-19, the vitamin stimulates the antiviral response induced by IFNs, regulates the exacerbated immune response induced by NF-kappaB and favors the inhibition of gene expression. Regarding the SEV group, the results indicate that in addition to suppressing viral infection, B12 induces inflammatory attenuation during severe COVID-19 and promotes activation of the adaptive immune response. These processes seem to have been regulated by epigenetic mechanisms, such as changes in chromatin status, gene silencing by miRNAs and alternative splicing reduction, which favored the reduction of gene transcription and translation.

Functional enrichment analysis also disclosed B12-modulated pathways in the MOD (Fig. 5B and Fig. S5) and/or SEV (Fig. 5C and Fig. S6) groups. It should be noted that the effect of B12 was pleiotropic, that is, it regulates several metabolic or intracellular signaling pathways maladjusted by COVID-19 despite the glucocorticoid treatment that patients received during hospitalization. Most of the differentially regulated pathways in contrasts 1 and 2 (MOD vs. CTRL and SEV vs. CTRL, all at endpoint A) had their activation or inhibition attenuated by B12 (contrasts 3 and 4 - MOD at endpoint B vs. CTRL at endpoint A; and SEV at endpoint B vs. CTRL at endpoint A). In the cultures of patients with moderate COVID-19, B12 favored the overall reduction of inflammation and activation of the Th1-type antiviral response (Fig. 5B and Fig. S5). The few pathways predicted to be more activated in response to B12, such as *iNOS signaling*, regulate desirable mechanisms in the context of COVID-19. Through S-nitrosylation of cysteine residues of viral and host proteins, NO reduces the activity of viral proteases, inhibiting the fusion and replication of SARS-CoV-2 in host cells (*46*). Another example is the *NK cell signaling pathway*. During infection, the coronavirus depletes NK cells and impairs their antiviral effects while activating macrophages and other immune cells leading to the cytokine storm (*47*). Furthermore, the activation of *TREM1 signaling* and *Acute phase response signaling* pathways favors the production of IFN and activation of Th1 and Th17 cells, which are fundamental components of the antiviral response (*32, 48, 49*). Activation of the *Th1 pathway* was also observed, which further reinforces the idea that B12 stimulates the antiviral response. Regarding the few pathways where the result seems incongruous, when analyzing their topology, it is clear that their activation by B12 is potentially beneficial to the patient. For example, in the *Inflammasome pathway*, NLRP1 was inhibited in the MOD group and B12 abrogates this inhibition (Fig. S8). This inflammasome interacts with double-stranded RNA (ds), being an important detector of dsRNA viruses such as SARS-CoV-2 (*50*). NLRP1 activity is regulated by anti-apoptotic proteins, BCL-2 and BCL-XL, which associate with the inflammasome and inhibit its activity (*51*). In this study, the genes encoding these two proteins were predicted to be upregulated in the leukocytes of patients in the MOD group, however B12 did not affect their expression. Thus, the regulatory effect of B12 on NLRP1 likely depends on other mediators. In the *Coronavirus Pathogenesis Pathway*, angiotensin 1-7 was predicted to be the activated in cultures treated with B12 (Fig. S7). It binds to MAS1 receptors leading to the activation of the anti-inflammatory branch of the renin-angiotensin pathway (*52*).

Moreover, in this same pathway, *Accumulation of lungs edema fluids* and *SARS CoV Replication* are predicted to be inhibited and *Type I interferon response* and *Adaptive Immunity* are predicted to be activated by B12. The vitamin caused even more evident attenuation of most of the inflammatory pathways activated by the disease in the cultures of patients with severe COVID-19 (Fig. 5C). In summary, the pro-inflammatory transcriptional profile observed in leukocytes from patients with moderate and severe COVID-19 was clearly attenuated by vitamin B12, with no adverse effects predicted from these results.

Besides regulating DNA methylation acting as a cofactor of the enzyme MS in the sulfur amino acid pathway, B12 also modulates different chromatin remodeling mechanisms that can affect gene expression. In the Krebs cycle, AdoCbl can induce the accumulation of succinate, an antagonist of histone and DNA demethylation reactions catalyzed by enzymes of the 2-oxoglutarate-dependent dioxygenase family, which may favor the activation or inhibition of gene expression (*53*). The antioxidant activity of B12 (*54*) can also indirectly influence the epigenetic landscape by regulating oxidative stress, which, depending on the context, can determine an accessible or non-accessible state of chromatin. Oxidative stress inactivates the histone-deacetylating enzyme HDAC2, contributing to the greater accessibility of transcriptional machinery to DNA. However, oxidative stress also activates signaling pathways that increase the expression of the enzyme DNMT1 that methylates DNA (*53, 55*). All these mechanisms may contribute to the down- (1,364) or upregulation (1,335) of the 2,699 genes in the leukocytes of patients in the MOD and/or SEV groups in response to B12 (Fig. 4).

It is noteworthy that when patients were recruited to this study, in July 2020, vaccines were not yet available and dexamethasone and other glucocorticoids had recently been integrated to the COVID-19 therapeutic protocol. At that time, there was no record of SARS-CoV-2 variants of interest/concern circulating in Brazil. Therefore, caution is needed when extrapolating the findings of this work to COVID-19 patients previously vaccinated or infected with SARS-CoV-2 variants.

In conclusion, vitamin B12 has a great potential as an adjuvant drug for alleviating inflammation in patients with moderate or severe COVID-19 in addition with the other established treatments. Beyond favorably regulating the expression of several inflammatory genes via methyl-dependent epigenetic mechanisms, B12 also acts as an antioxidant, has an excellent safety profile and is widely available at low cost. Therefore, phase II/III clinical trials are enthusiastically recommended. Furthermore, the results presented herein neither endorse the prophylactic use of B12 to prevent SARS-CoV-2 infection, nor its therapeutic use in mild COVID-19.

## Materials and Methods

### Patients

This study was approved by the National Research Ethics Committee of Brazil (CONEP: 32242620.6.0000.5091). All patients or their legal representatives were informed about the study and signed an informed consent form. Patients with moderate (MOD; n = 10) and severe (SEV; n = 16) forms of COVID-19, classified according to the World Health Organization’s COVID-19 clinical severity scale (*56*), admitted to the Metropolitan Hospital Doctor Célio de Castro (HMDCC) and uninfected volunteers (CTRL; n = 6) with biological sex and age parity with the patients were recruited from 20th to 26th July 2020. Participants had peripheral venous blood samples (10 mL) collected in sodium heparin, at 8 a.m., and data from medical records of COVID-19 patients were analyzed. Uninfected volunteers included as controls in the study were tested for the presence of SARS-CoV2 in the oropharynx and nasopharynx by RT-qPCR and for the presence of IgM and IgG blood antibodies against the virus by lateral flow immunochromatography (Wondfo, Guangzhou, China).

### Sample processing and production of whole blood cultures

Each peripheral venous blood sample was divided into 4 aliquots of 1 mL and processed as follows: endpoint U) unprocessed blood; endpoints A and Z) added with 498 µl of RPMI 1640 culture medium (Sigma-Aldrich, Saint Louis, Missouri) and 2 µl of pH 5 citrate-phosphate buffer excipient (Merck, Darmstadt, Germany); and endpoint B) added with 500 µl of RPMI 1640 culture medium with cyanocobalamin (Merck) to a final concentration of 1 nM. Aliquots Z were immediately processed with no incubation. Aliquots A and B were incubated in 6-well plates for 24 hours at 37°C in a humidified atmosphere with 5% CO_2_ (Fig. 6).

**Fig. 6.**
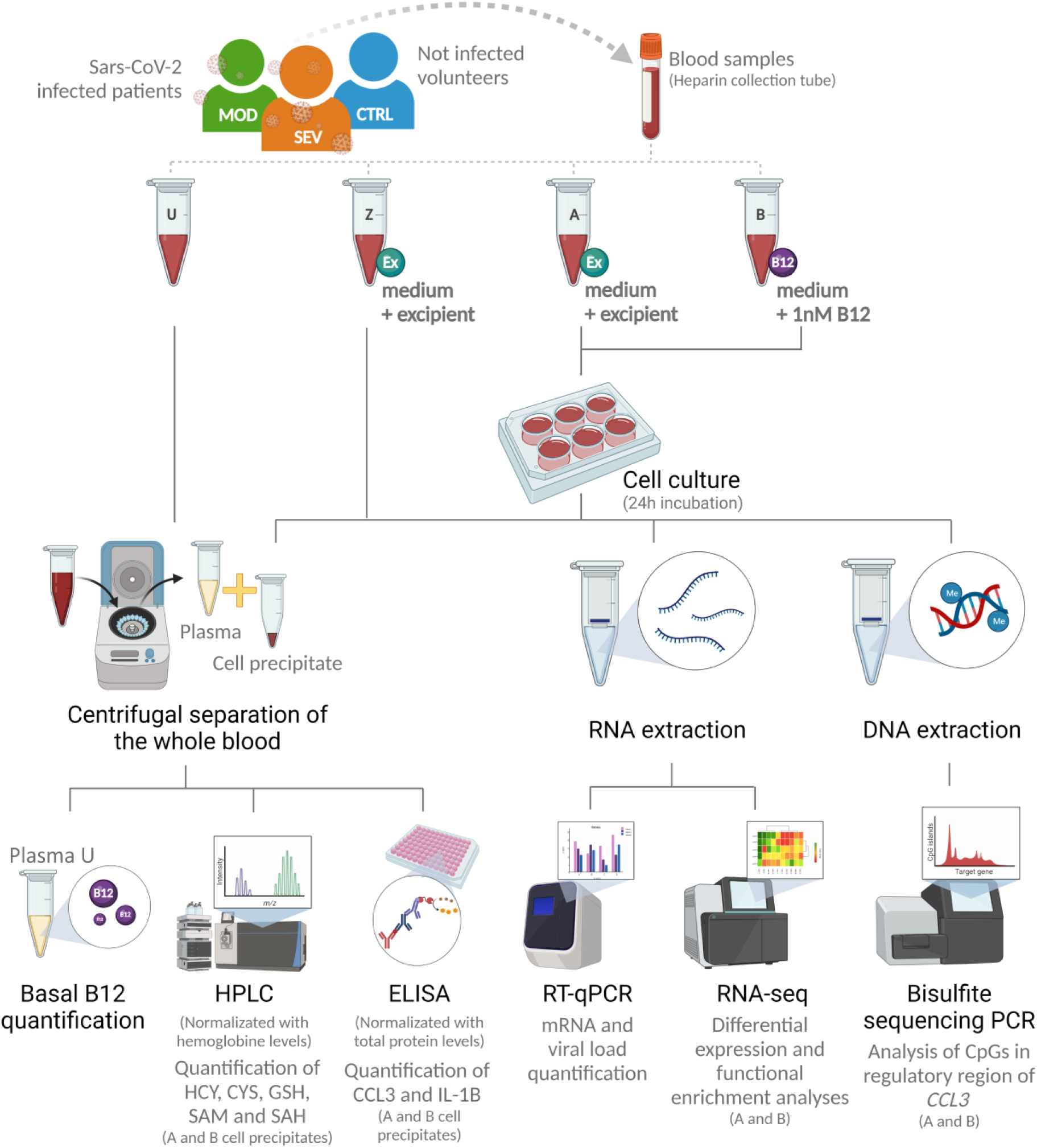
Research design. Abbreviations: SEV = severe COVID-19; MOD = moderate COVID-19; CTRL = non-infected controls; U = endpoint U (unprocessed blood); Z = endpoint Z (samples immediately processed after addition of culture medium with excipient); A = endpoint A (Samples added to culture medium with excipient and incubated for 24h); B = endpoint B (Samples added to culture medium with B12 and incubated for 24h); Ex = Excipient; B12 = Vitamin B12; HPLC = High Performance Liquid Chromatography; HCY = Homocysteine; CYS = Cysteine; GSH = Glutathione; SAM = S-adenosylmethionine; SAH = S-adenosyl-L-homocysteine; PCR = Polymerase Chain Reaction; RT-qPCR = Real-time quantitative PCR. (Created with BioRender.com).

### Quantification of basal B12

Basal levels of vitamin B12 were quantified of plasma aliquots from endpoint U using the standard chemiluminescence method by a commercial clinical laboratory (Hermes Pardini, Belo Horizonte, Brazil).

### Real time quantitative PCR (RT-qPCR)

Total RNA was obtained from 1.5 mL of aliquots from endpoints Z, A and B, using the QIAamp RNA Blood kit (Qiagen, Hilden, Germany), and cDNA synthesis was performed from 2 µg of total RNA using the High-Capacity cDNA Reverse Transcription Kit (ThermoFisher, Waltham, Massachusetts), both according to manufacturers’ protocols. All samples had their RNA quantified by fluorometry using the Qubit RNA HS Assay Kit (#Q32852) and the Qubit 2.0 fluorometer (Invitrogen, Carlsbad, CA).

Specific primers (Table S2) were used to detect the human mRNAs of *CCL1* (C-C Motif Chemokine Ligand 1) (NM_002981.2), *CCL2* (C-C Motif Chemokine Ligand 2) (NM_002982.4), *CCL3* (C-C Motif Chemokine Ligand 3) (NM_002983.3), *CXCL9* (C-X-C motif chemokine ligand 9) (NM_002416.3), *IL1B* (Interleukin 1 beta) (NM_000576.3), *IL6* (Interleukin 6) (NM_000600.5), *IL17A* (Interleukin 17A) (NM_002190.3), *TNF* (Tumor Necrosis Factor) (NM_000594.4), *HAVCR2* (Hepatitis A Virus Cellular Receptor 2) (NM_032782.5), *CD4* (CD4) (NM_000616.5) and *CD8A* (CD8a) (NM_001768.7). For all RT-qPCR assays, target gene expression levels were normalized by 18S ribosomal RNA (18S ribosomal N1) (NR_145820.1) levels. qPCR reactions were performed with Fast SYBR Green Master Mix (ThermoFisher) plus 10 ng of cDNA in a final volume of 10 µL. Thermal cycling and fluorescence detection were performed using the ViiA 7 real-time PCR system (ThermoFisher) according to the manufacturer’s recommendations. The relative expression of target genes was calculated using the 2e(-ΔCt) method (*57*).

### ELISA

To access the intracellular protein levels of CCL3 and IL-1B, cell precipitates of aliquots from endpoints A and B were homogenized in 1X PBS, diluted in Milli-Q H_2_O 1:2 and the cells disrupted by mechanical lysis in the TissueLyser (Qiagen) for 2 minutes at 30 Hz and three freeze/thaw cycles (20° to -20°C).

ELISA kits (R&D Systems, Abingdon, UK) for CCL3 (#DY270) and IL-1B (#DY201) were used and assays performed as per the manufacturer’s instructions. The absorbance was read at 450 nm with the Multiskan GO spectrophotometer (ThermoFisher). Standard curves were constructed from 0.1 to 100 pg/mL and cytokine concentrations were calculated with SkanIt Software 4.1 for Microplate Readers RE, version 4.1.0.43 (ThermoFisher).

### High performance liquid chromatography (HPLC)

For measurements of HCY, cysteine (CYS), GSH, SAM and S-adenosyl-L-homocysteine (SAH), cell precipitates from aliquots A and B were diluted in Milli-q H_2_O 1:2 and the cells disrupted by three freeze/thaw cycles (20° to -20°C). Metabolites were quantified by high performance liquid chromatography (HPLC) on the Shimadzu Prominence-i LC-2030C 3D Plus (GMI, Ramsey, Minnesota) according to Pfeiffer *et al*. (*58*) with some modifications. HCY, CYS and GSH were quantified using a C18 Luna column (5 mm x 150 mm x 4.6 mm) and mobile phase composed of 0.06 M sodium acetate, 0.5% acetic acid and 2% methanol (pH 4.7 adjusted with acetic acid). The flow rate was 1.1 mL × min^1^ and the retention time was 4.1 min for CYS, 5.9 min for HCY and 10.3 min for GSH. SAM and SAH were quantified using a method adapted from Blaise *et al*. (*59*). HClO_4_ was added to the homogenized cell precipitate for protein precipitation. The supernatant was injected into a C18 LiChroCart column (5 mm x 250 mm x 4 mm) and the mobile phase applied at a flow rate of 1 mL × min^− 1^, consisting of 50 mM sodium phosphate (pH 2.8), 10 mM heptane sulfonate and 10% acetonitrile. Retention time was 8.7 min for SAH and 13.6 min for SAM. All metabolites were detected by UV absorption at a wavelength of 254 nm.

The intracellular concentrations of the metabolites were normalized by hemoglobin concentrations, obtained by colorimetry (Agabe hemoglobinometer) adding 10 µL of the sample in an ampoule (Hemoglobin AP; Labtest).

### RNA-Seq

Total RNA from three samples from each group (MOD, SEV and CTRL) at endpoints A and B (total 18 samples) was used to construct cDNA libraries with the TruSeq Stranded mRNA kit (Illumina, San Diego, CA) and the indexed fragments were sequenced on the NGS NextSeq 500 (Illumina) with the NextSeq 500/550 High Output 2×75 cycles kit (Illumina). All sequenced samples were quality assessed by capillary electrophoresis with the Agilent RNA 2100 Nano kit (Bioanalyzer, Santa Clara, CA).

### Bisulfite Sequencing PCR (BSP)

Genomic DNA from five samples from each group (MOD, SEV and CTRL) at endpoints A and B (total of 30 samples) was extracted with QIAamp DNA Blood Mini kit (Qiagen) according to the manufacturer’s instructions. DNA was subjected to bisulfite conversion using EpiTect Bisulfite kit (Qiagen). The region -107pb to +164pb of *CCL3* (GRCh38/hg38 Chr17: 36,090,276-36,090,005) that cover the promoter region and proximal portion of the first exon of the gene was PCR amplified with GoTaq DNA Polymerase (Promega, Madison, Wisconsin) from bisulfite-converted DNA using the primer listed in Table S3. DNA libraries were purified with AMPure XP beads (Beckman Coulter, Indianapolis, IN), indexed with Nextera XT DNA Library Preparation Kit (Illumina) and sequenced on the NGS MiSeq (Illumina) with MiSeq Reagent kit v2 (300 cycles) (Illumina). All sequenced samples had their quality evaluated by capillary electrophoresis with Agilent High Sensitivity DNA Kit (Bioanalyzer).

### Bioinformatics analysis

Hierarchical clustering analyzes were performed using the GenePattern software (*60*) using Pearson correlation as a comparison method and average linkage as a linkage method.

Raw RNA-Seq reads were pre-processed with Trimmomatic software (*61*) to remove adapters, poor quality bases or very short reads (less than 36nt). The filtered reads were mapped to the *Homo sapiens* reference transcriptome (Gencode, release 36) and the total number of reads mapped per transcript and the number of transcripts per million (TPM) were calculated with Salmon software (*62*). Transcript counts and abundances were summarized at the gene level using the *summarizeToGene* function of the R Tximeta package (*63*). Contrast analyses between MOD vs. CTRL at endpoint A (contrast 1), SEV vs. CTRL at endpoint A (contrast 2), MOD at endpoint B vs. CTRL at endpoint A (contrast 3), SEV at endpoint B vs. CTRL at endpoint A (contrast 4), and CTRL at endpoint A vs. CTRL at endpoint B (contrast 5) were performed with the R DESeq2 package (*64*). Genes with Fold Change greater than 1.5 and *P* value less than 0.05 obtained in the False Discovery Rate (FDR) analysis were considered differentially expressed (DEG) and submitted to functional enrichment analysis with Ingenuity Pathways Analysis software (IPA, Qiagen) using default parameters. Pathways with -log *P* value greater than 2 were considered to be differentially activated or inhibited. To assess the differences between moderate and severe COVID-19 (contrast 1 vs. contrast 2) and the effect of B12 (contrast 3 vs. contrast 1, and contrast 4 vs. contrast 2), only differentially regulated pathways that had more than 20% difference in Z-scores between the two contrasts were selected. Genes differentially expressed in response to B12 were submitted to gene ontology (GO) functional annotation analysis using *DAVID Bioinformatics* software (*65*) with default parameters. The GO terms for Biological Processes with *P* < 0.05 were summarized according to their ontology with REVIGO (*66*).

For BSP analysis, raw reads were preprocessed with Trimmomatic (*61*) as described above, except that the minimum read size threshold was 50nt. Since no single nucleotide polymorphisms (SNPs) that influence CpGs dinucleotides, which can generate or abolish a CpG site, were identified in the analyzed region, the reference sequence chr17: 36,090,276-36,090,005 of the genome of *Homo sapiens* (Gencode, release 38), was used to map the filtered reads. The mapping, as well as the calculation of the percentages of methylation of cytosines in CG context were performed with the Bismark software (*67*). *CCL3* functional annotations, such as transcription start site (TSS), SNPs and predicted transcription factor binding sites (TFBS) were identified with the UCSC Genome Browser (*68*).

### Statistical analysis

Statistical analyses were performed using GraphPad Prism software (version 8.0.2) (GraphPad Software Inc., Irvine, CA). Data distribution was analyzed using Anderson-Darling, D’Agostino & Person, Shapiro-Wilk and Kolmogorov-Smirnov tests. For comparisons between three or more groups, one-way analysis of variance (ANOVA) or Kruskall-Wallis tests were used, followed by the multiple comparison tests of Tukey or Dunn, for parametric or non-parametric data, respectively. Two-way ANOVA followed by Tukey’s multiple comparison test was used for comparisons involving the effect of two factors on a dependent variable in three or more groups. Two-tailed Student’s t, paired Student’s t, Mann-Whitney or Wilcoxon tests were used for comparisons between two groups, according to the experimental design and data distribution. Correlations were tested using Pearson or Spearman tests according to data distribution. Outliers identified by ROUT test (Q = 1%) were excluded. Data were expressed as median ± interquartile (nonparametric) or mean ± standard deviation (parametric). Differences were considered statistically significant when *P* < 0,05.

For BSB analysis, the means of methylation percentages of each CG locus of MOD, SEV and CTRL groups at endpoints A and B were compared using two-tailed Wilcoxon test and only those that presented a statistically significant difference were considered. Correlation between methylation levels of differentially methylated loci (DML) at endpoint B compared to endpoint A and *CCL3* gene expression levels from the same cultures were tested with Pearson or Spearman correlation for parametric or nonparametric distributions, respectively.

## Acknowledgments

This work received financial support from FIOCRUZ, CAPES, INCT-Vacinas and from a gentle personal money donation by Claudia Garcia Martins. Technical support was provided by FIOCRUZ Technological Platforms: Real Time PCR and Digital PCR (P04-006), NGS Sequencing (P01-007), and Bioinformatics (P08-002). We thank Dr. Ludmila R.P. Ferreira for valuable discussions about the functional enrichment analysis, and Dr. Diana Bahia and Prof. Dulciene M.M. Queiroz for critically reviewing this manuscript.

## Author contributions

LC - performed the experiments, contributed to experimental design, analyzed data and wrote the manuscript; VCS and VD’A – performed the HPLC assays, analyzed data and reviewed the manuscript; CC, BP and SF – recruited patients, collected blood samples and clinical and sociodemographic data from patients records; MO - contributed to data analysis and reviewed the manuscript; AS - made substantial contributions to the NGS experiments; GF - contributed to the experimental design and data analysis and reviewed the manuscript; RC - designed and oversaw the study, analyzed data and wrote the manuscript. All authors read and approved the final manuscript.

## Competing interests

The authors declare that they have no competing interests.

## Data and materials availability

The dataset supporting the conclusions of this article is available in the SRA repository, [https://www.ncbi.nlm.nih.gov/sra/PRJNA862565]. Other data needed to evaluate the conclusions in the paper are present in the Supplementary Materials.

## Supplementary Materials

Fig. S1. Vitamin B12 basal levels

Fig. S2. Samples added to culture medium with excipient and incubated for 24h have their transcriptional signatures preserved

Fig. S3. Biological Processes affected by B12

Fig. S4. Patients with moderate and severe COVID-19 previously treated with glucocorticoids had distinct global gene expression patterns

Fig. S5. Vitamin B12 attenuated the pro-inflammatory profile of leukocytes from patients with moderate COVID-19

Fig. S6. Vitamin B12 attenuated the pro-inflammatory profile of leukocytes from patients with moderate COVID-19

Fig. S7. Coronavirus Pathogenesis Pathway (Contrast 1, 2, 3 and 4) Fig. S8. Inflammasome Pathway (Contrast 1 and 3)

Table S1. Patients’ data (separate file)

Table S2. COVID-19 hyperinflammation-related genes panel primers

Table S3. BSP primers

## Notes

### Competing Interest Statement

The authors have declared no competing interest.

