## Supplementary Materials for "Vitamin B12 attenuates leukocyte inflammatory signature in COVID-19 via methyl-dependent changes in epigenetic marks"

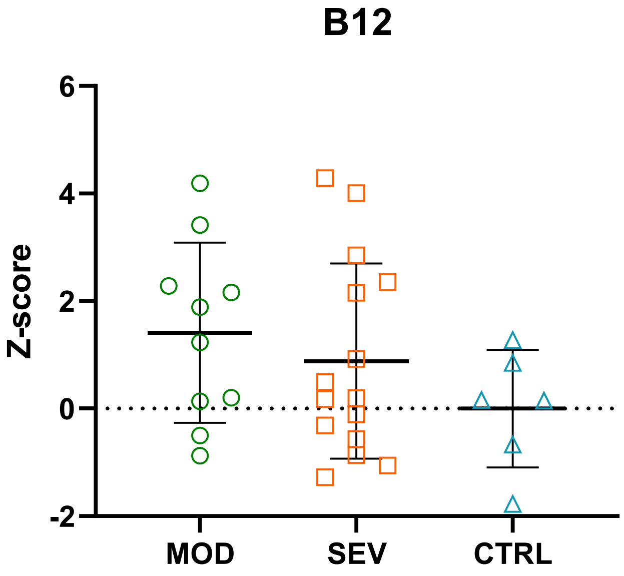


**Fig. S1. Vitamin B12 basal levels.** Plasma levels (endpoint Br) of vitamin B12 in patients with moderate (MOD) or severe (SEV) COVID-19 and non-infected controls (CTRL). Dosages were compared using the one-way Analysis of Variance (ANOVA) test followed by Tukey's multiple comparisons test and expressed as mean ± standard deviations of Z-score values. N sample = MOD (10), SEV (16), CTRL (6).

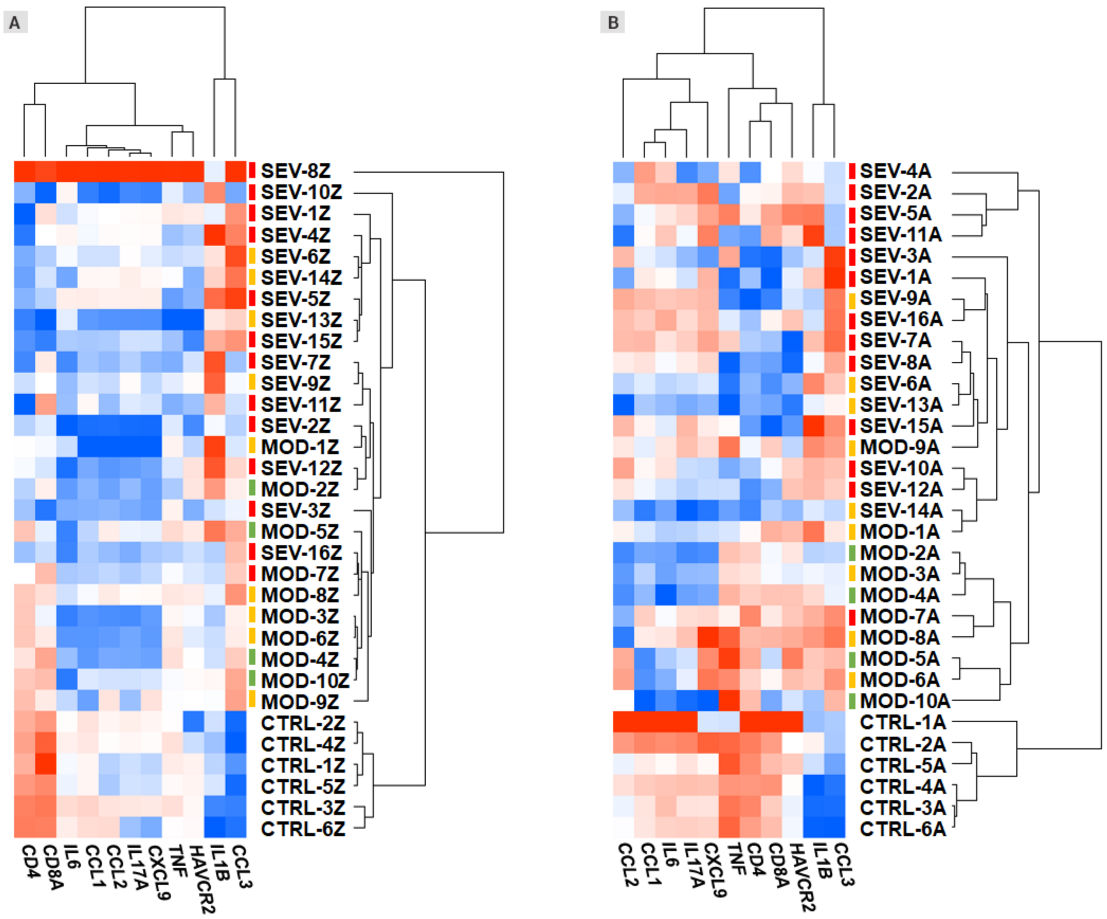


**Fig. S2. Samples added to culture medium with excipient and incubated for 24h have their transcriptional signatures preserved. A** and **B**: Dendrogram and heatmap depicting the hierarchical clustering and expression levels of COVID-19 hyperinflammation-related genes panel in whole blood cultures of infected patients (MOD and SEV) and non-infected controls at endpoint Z (Panel A) and endpoint A (Panel B). The 2e(-ΔCt) values for gene expression levels were grouped by hierarchical clustering parameterized with Pearson correlation and average linkage. Heatmap colors were normalized by column. Colored dashes next to the sample identification correspond to the patient's outcome. Green dashes: patients discharged from hospital one day after sample collection; Yellow lines: patients discharged from hospital two or more days after sample collection; Red lines: patients' death. Abbreviations: SEV = severe COVID-19; MOD = moderate COVID-19; CTRL = non-infected controls. Suffixes: Z = endpoint Z (samples immediately processed after addition of culture medium with excipient); A = endpoint A (samples added to culture medium with excipient and incubated for 24h).

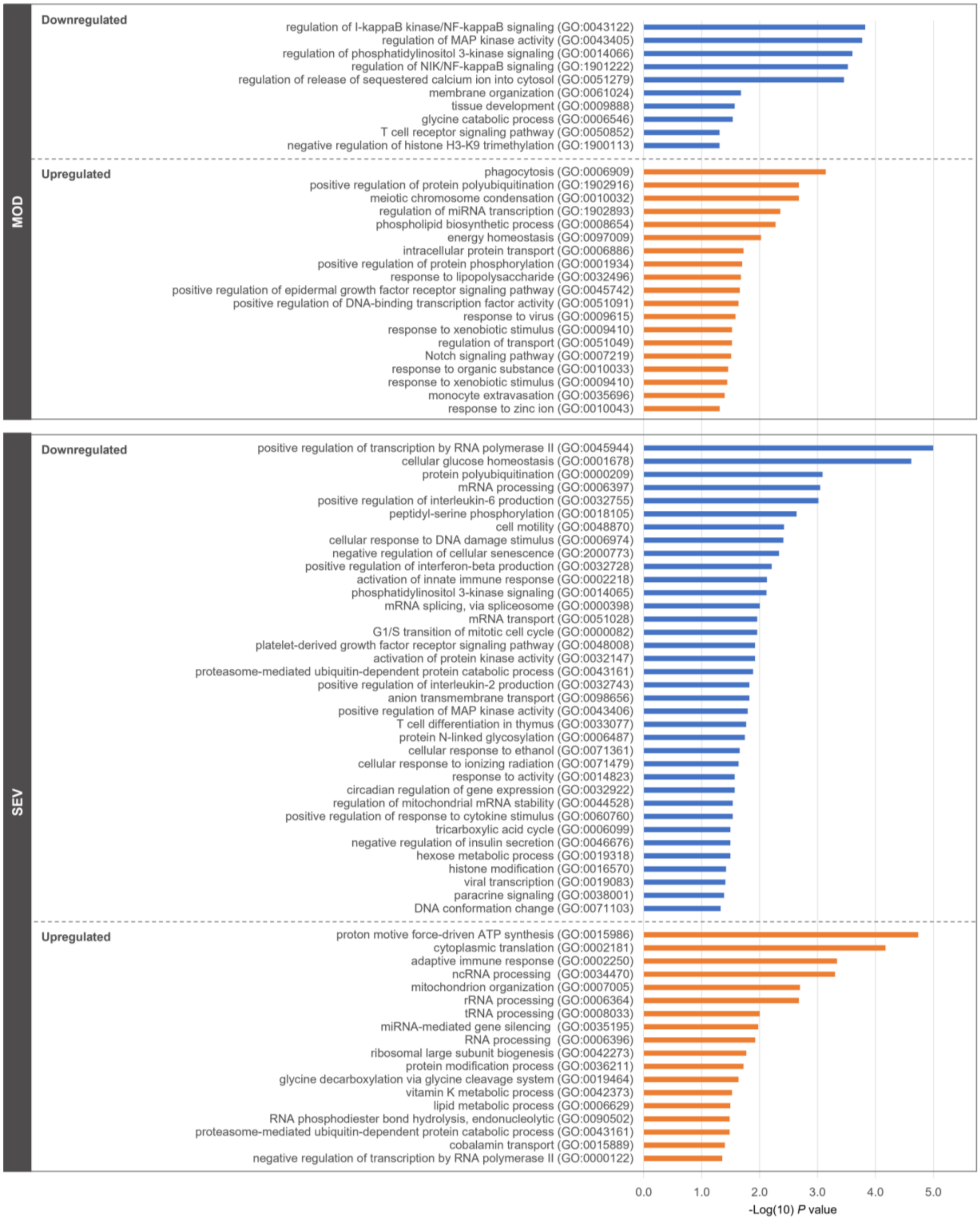


**Fig. S3. Biological Processes affected by B12.** GO terms for Biological Processes with $P < 0.05$. Abbreviations: SEV = severe COVID-19; MOD = moderate COVID-19.

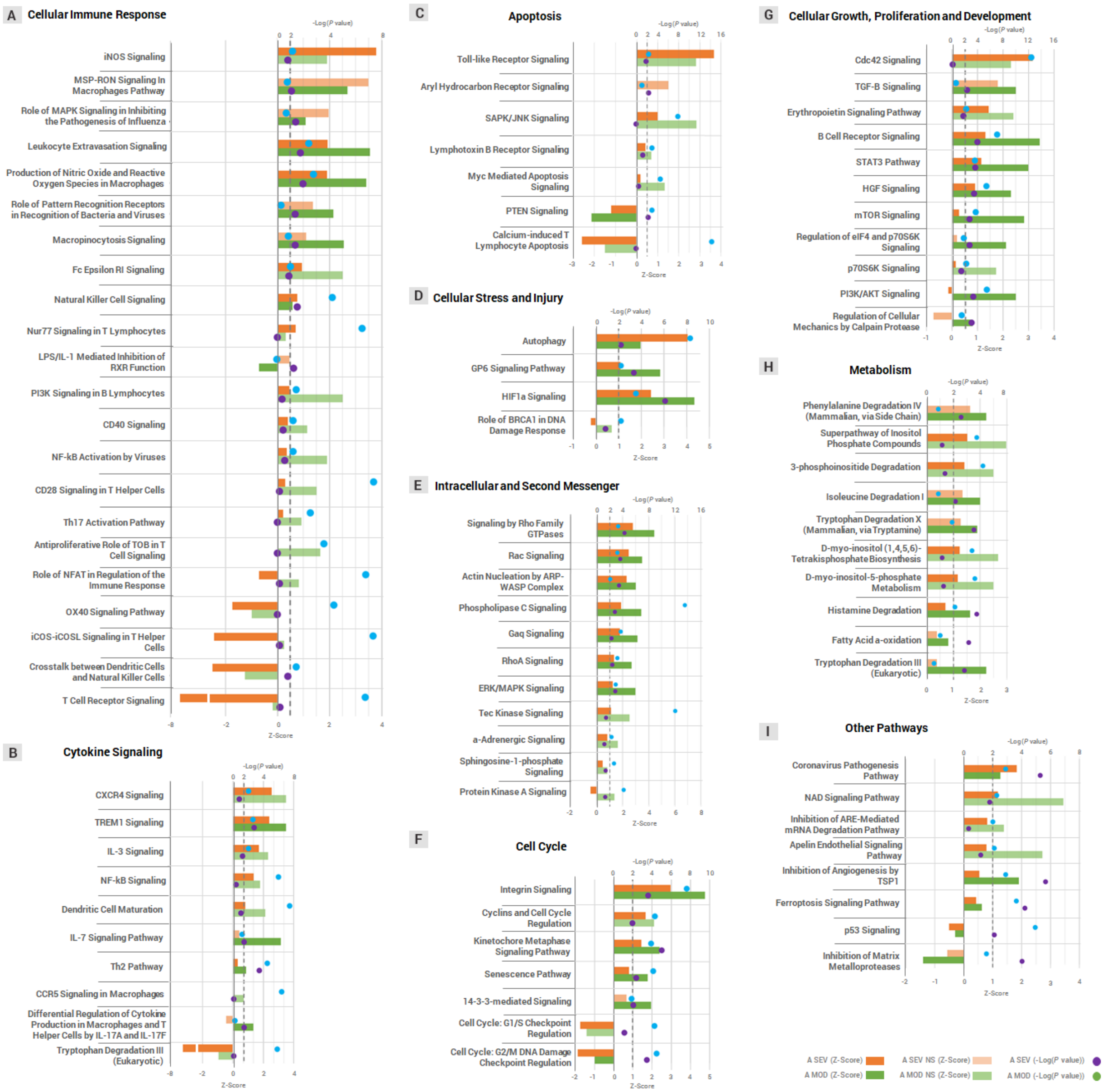


**Fig. S4. Patients with moderate and severe COVID-19 previously treated with glucocorticoids had distinct global gene expression patterns.** Canonical pathways differentially regulated that had more than 20% difference in Z-scores between contrast 1 and 2. Abbreviations: NS = statistically non-significant; SEV = severe COVID-19; MOD = moderate COVID-19. Suffixes: A = endpoint A (Samples added to culture medium with excipient and incubated for 24h).

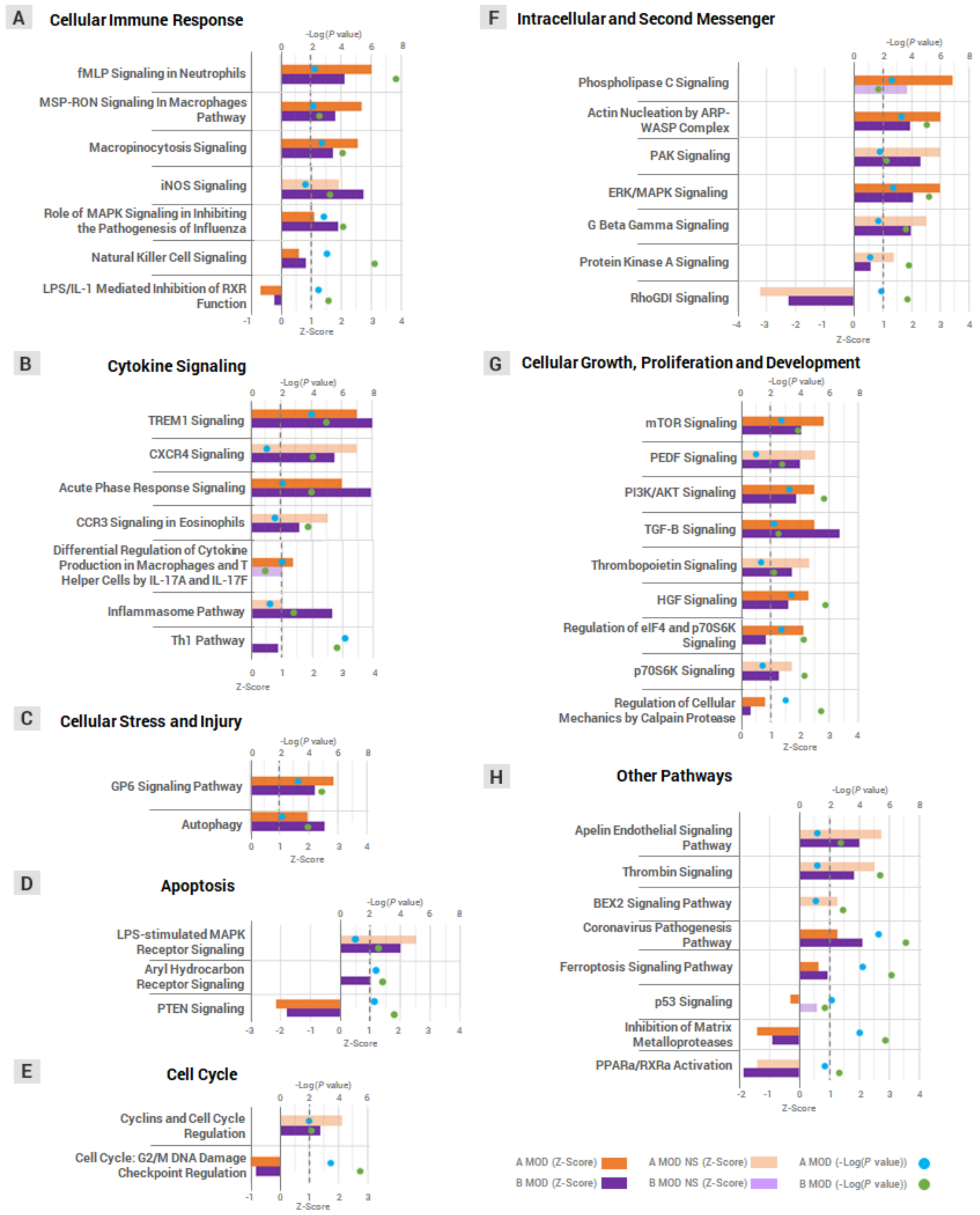


**Fig. S5. Vitamin B12 attenuated the pro-inflammatory profile of leukocytes from patients with moderate COVID-19.** Canonical pathways differentially regulated that had more than 20% difference in Z-scores between contrast 1 and 3. Abbreviations: NS = statistically non-significant; MOD = moderate COVID-19. Suffixes: A = endpoint A (Samples added to culture medium with excipient and incubated for 24h); B = endpoint B (Samples added to culture medium with B12 and incubated for 24h).

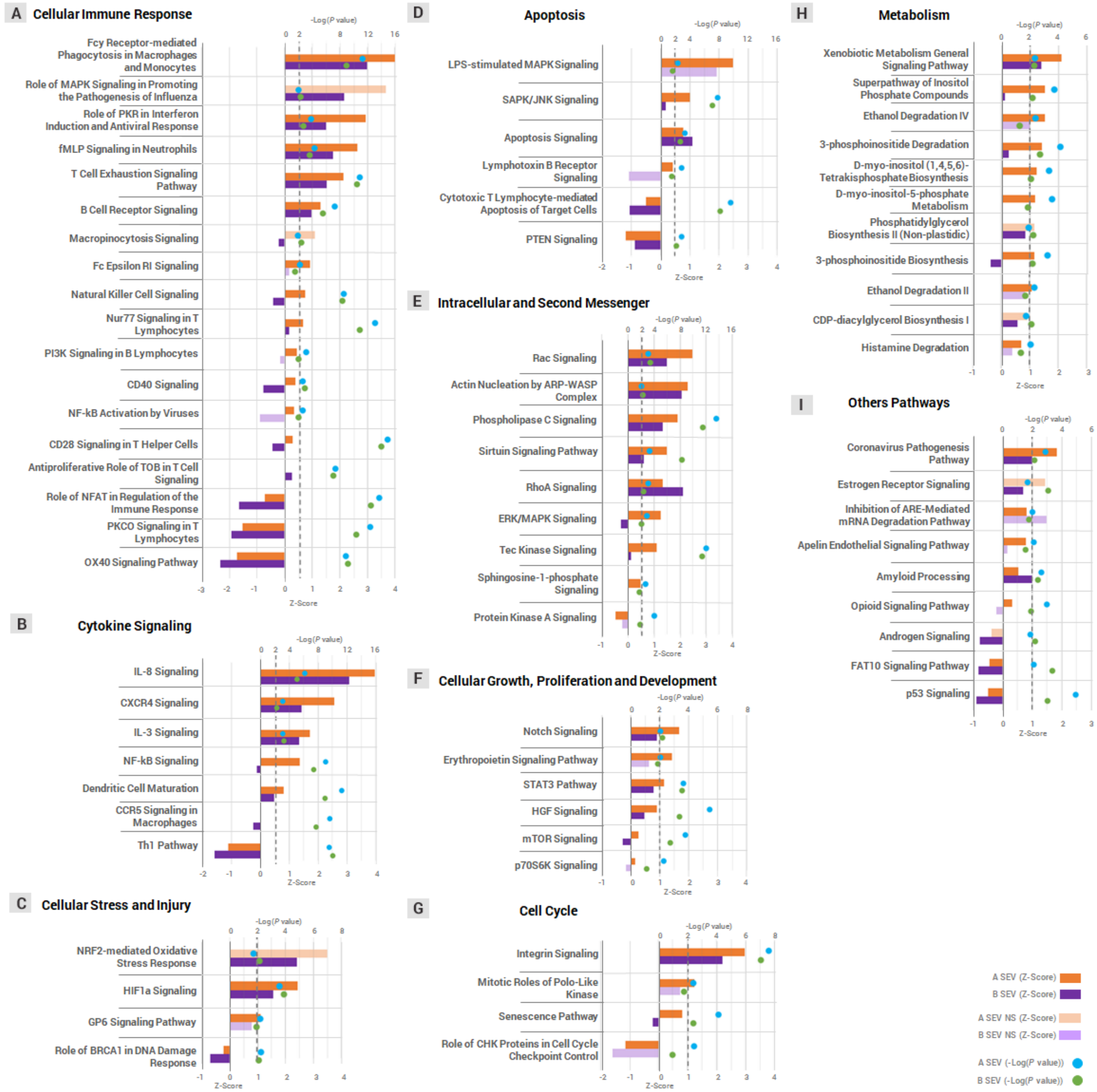


**Fig. S6. Vitamin B12 attenuated the pro-inflammatory profile of leukocytes from patients with severe COVID-19.** Canonical pathways differentially regulated that had more than 20% difference in Z-scores between contrast 1 and 3. Abbreviations: NS = statistically non-significant; SEV = severe COVID-19. Suffixes: A = endpoint A (Samples added to culture medium with excipient and incubated for 24h); B = endpoint B (Samples added to culture medium with B12 and incubated for 24h).

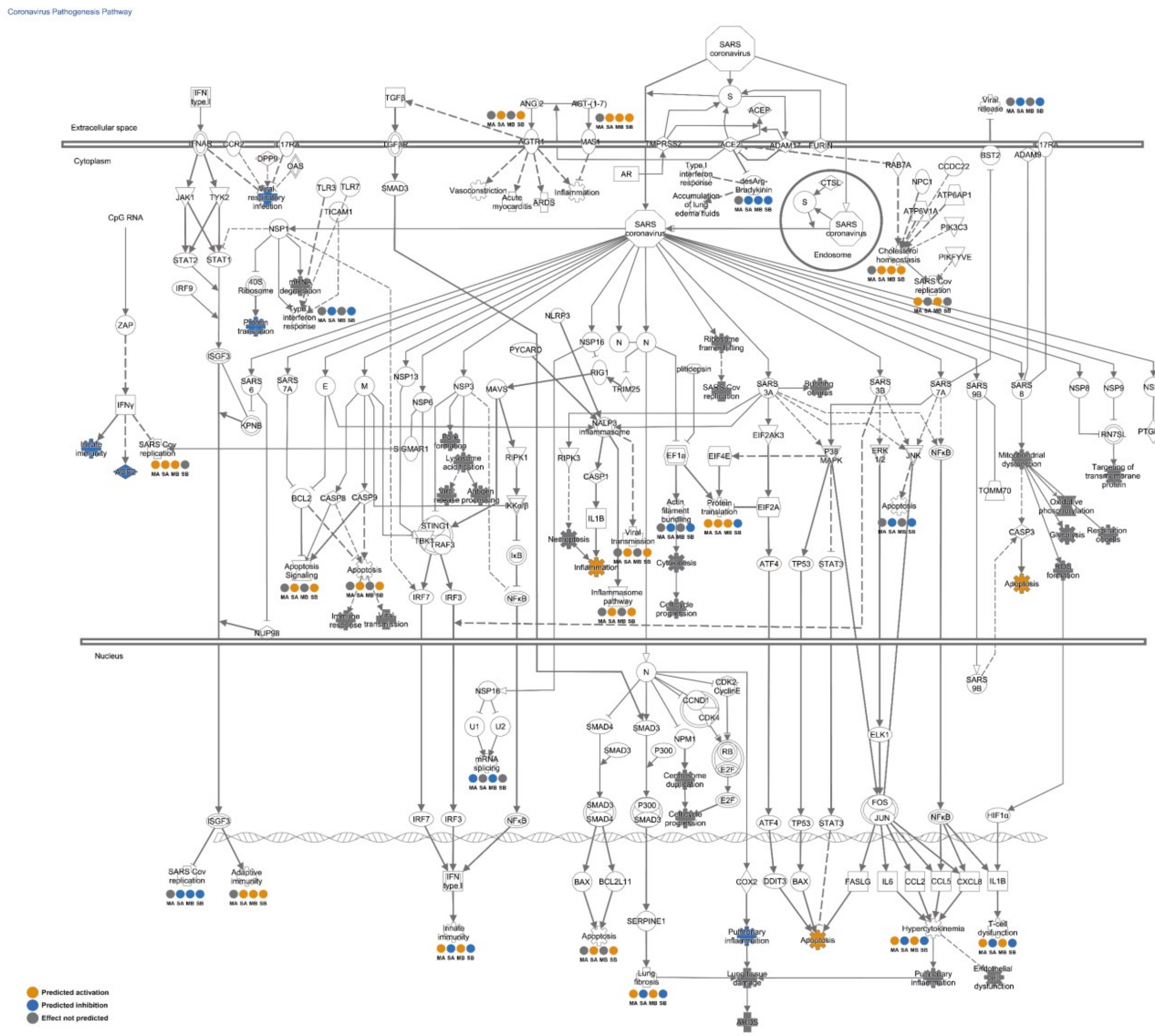


**Fig. S7. Coronavirus Pathogenesis Pathway (Contrast 1, 2, 3 and 4).** Abbreviations: MA = moderate COVID-19 at endpoint A (contrast 1); MB = moderate COVID-19 at endpoint B (contrast 3); SA = severe COVID-19 at endpoint A (contrast 2); SB = severe COVID-19 at endpoint B (contrast 4); A = endpoint A (samples added to culture medium with excipient and incubated for 24h); B = endpoint B (samples added to culture medium with B12 and incubated for 24h); up = upregulated; down = downregulated. Contrast 1 = MOD vs. CTRL at endpoint A; Contrast 2 =SEV vs. CTRL at endpoint A; Contrast 3 = MOD at endpoint B vs. CTRL at endpoint A; and Contrast 4 = SEV at endpoint B vs. CTRL at endpoint A. Adapted from Ingenuity Pathways Analysis (IPA, Qiagen).

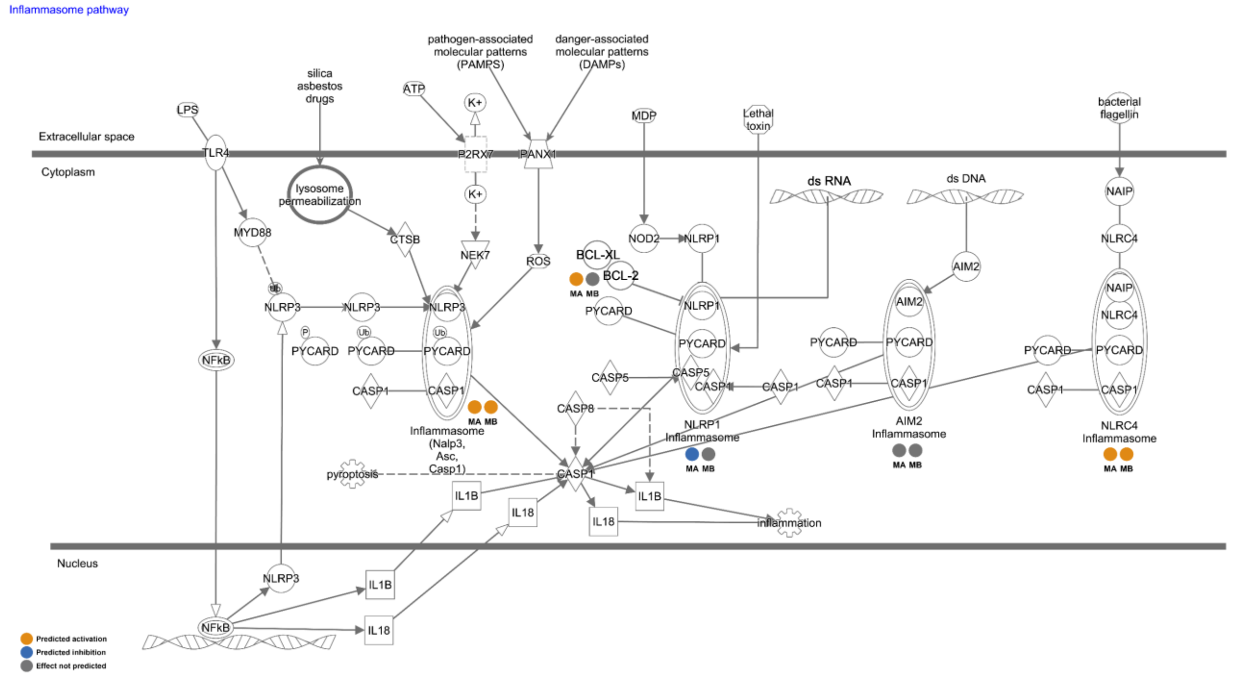


**Fig. S8. Inflammasome Pathway (Contrast 1 and 3).** Abbreviations: MA = moderate COVID-19 at endpoint A (contrast 1); MB = moderate COVID-19 at endpoint B (contrast 3); A = endpoint A (samples added to culture medium with excipient and incubated for 24h); B = endpoint B (samples added to culture medium with B12 and incubated for 24h); up = upregulated; down = downregulated. Contrast 1 = MOD vs. CTRL at endpoint A; and Contrast 3 = MOD at endpoint B vs. CTRL at endpoint A. Adapted from Ingenuity Pathways Analysis (IPA, Qiagen).

| Information | | All (n=26) | MOD (n=10) | SEV (n=16) | *P* value (MOD x SEV) |
|---|---|---|---|---|---|
| **General Information** | | | | | |
| **Biological Sex** | Female [n (%)] | 14 (55.56) | 5 (54.55) | 9 (56.25) | >0.9999 |
| | Male [n (%)] | 12 (44.44) | 5 (45.45) | 7 (43.75) | |
| **Age [years; median (IQ)]** | | 64 (59.75-71) | 64 (58.5-72.75) | 63.5 (59.5-69.5) | 0.9341 |
| **Blood type [n (%)]** | | O+ [8 (30.77)] | O- [3 (30)] | O+ [6 (37.5)] | - |
| **Length of stay until the date of blood collection [days; mean (SD)]** | | 11.81 (4.205) | 11.10 (3.872) | 12.25 (4.465) | 0.5087 |
| **Time elapsed between symptom manifestation and admission to HMDCC [days; mean (SD)]** | | 6.333 (3.460) | 6.778 (4.116) | 6.067 (3.127) | 0.6365 |
| **Time elapsed between COVID test to blood draw [days; mean (SD)]** | | 5.423 (4.272) | 5.4 (2.547) | 5.438 (5.151) | 0.9832 |
| **Dexamethasone treatment [days; median (IQ)]** | | 11 (6.5-11.5) | 11 (6-11.5) | 10.5 (8.5-11.75) | 0.6546 |
| **Use of two or more glucocorticoides (Hydrocortisone and/or beclomethasone) [n (%)]** | | 14 (53.85) | 4 (40) | 10 (62.5) | 0.4216 |
| **Coexisting Medical Conditions** | | | | | |
| **Ex-Smokers [n (%)]** | | 5 (19.23) | 3 (30) | 2 (12.5) | 0.6241 |
| **Continuous use of Metformin [n (%)]** | | 4 (15.38) | 1 (10) | 3 (18.75) | 0.5865 |
| **Diabetes Mellitus [n (%)]** | | 10 (38.46) | 3 (30) | 7 (43.75) | 0.6834 |
| **Dyslipidemia [n (%)]** | | 2 (7.69) | - | 2 (12.5) | 0.5077 |
| **Obesity [n (%)]** | | 3 (11.54) | - | 3 (18.75) | 0.2615 |
| **Cardiovascular Diseases [n (%)]** | | 19 (73.08) | 8 (80) | 11 (68.75) | 0.668 |
| **Neurological Conditions [n (%)]** | | 5 (19.23) | 4 (40) | 1 (6.25) | 0.0549 |
| **Respiratory Diseases [n (%)]** | | 3 (11.54) | 1 (10) | 2 (12.5) | >0.9999 |
| **Kidney Diseases [n (%)]** | | 2 (7.69) | - | 2 (12.5) | 0.5077 |
| **Other Conditions [n (%)]** | | 4 (15.38) | 2 (20) | 2 (12.5) | 0.6254 |
| **Denies pre-existing condition [n (%)]** | | 5 (19.23) | 1 (10) | 4 (25) | 0.6169 |
| **Symptoms** | | | | | |
| **Cough [n (%)]** | | 18 (69.23) | 7 (70) | 11 (68.75) | >0.9999 |
| **Dispneia [n (%)]** | | 18 (69.23) | 6 (60) | 12 (75) | 0.6645 |
| **Fever [n (%)]** | | 12 (46.15) | 4 (40) | 8 (50) | 0.7015 |
| **Desaturation/ Hypoxemia [n (%)]** | | 10 (38.46) | 2 (20) | 8 (50) | 0.2177 |
| **Myalgia [n (%)]** | | 5 (19.23) | 2 (20) | 3 (18.75) | >0.9999 |

| | | | | |
|---|---|---|---|---|
| **Anosmia [n (%)]** | 3 (11.54) | - | 3 (18.75) | 0.2615 |
| **Loss of Appetite/Hyporexia/Inapetence [n (%)]** | 3 (11.54) | 2 (20) | 1 (6.25) | 0.5385 |
| **Diarrhoea [n (%)]** | 3 (11.54) | 2 (20) | 1 (6.25) | 0.5385 |
| **Nausea [n (%)]** | 2 (7.69) | 1 (10) | 1 (6.25) | >0.9999 |
| **Malaise [n (%)]** | 2 (7.69) | - | 2 (12.5) | 0.5077 |
| **Headache [n (%)]** | 2 (7.69) | 2 (20) | - | 0.1385 |
| **Dysgeusia [n (%)]** | 2 (7.69) | 1 (10) | 1 (6.25) | >0.9999 |
| **Rhinorrhea/ Runny Nose [n (%)]** | 2 (7.69) | 2 (20) | - | 0.1385 |
| **Sore Throat [n (%)]** | 2 (7.69) | - | 2 (12.5) | 0.5077 |
| **Chest Pain [n (%)]** | 2 (7.69) | 1 (10) | 1 (6.25) | >0.9999 |
| **Dizziness [n (%)]** | 2 (7.69) | 2 (20) | - | 0.1385 |
| **Prostration [n (%)]** | 1 (3.85) | - | 1 (6.25) | >0.9999 |
| **Mental Confusion [n (%)]** | 1 (3.85) | 1 (10) | - | 0.3846 |
| **Chills [n (%)]** | 1 (3.85) | - | 1 (6.25) | >0.9999 |
| **Flu syndrome [n (%)]** | 1 (3.85) | - | 1 (6.25) | >0.9999 |
| **Laboratory Findings - Biochemistry and Blood Count** | | | | |
| **Glucose [mg/dL; mean (SD)]** | 173.3 (63.83) | 131.7 (58.85) | 196.7 (55.2) | 0.0111 (*) |
| **Calcium [mg/dL; mean (SD)]** | 4.846 (0.389) | 4.732 (0.1811) | 4.918 (0.4673) | 0.2445 |
| **Lactate [mmol/L; median (IQ)]** | 2 (1.675-2.525) | 2.1 (1.75-2.725) | 1.85 (1.625-2.475) | 0.3546 |
| **Sodium [mmol/L; mean (SD)]** | 140.6 (6.645) | 139.8 (3.747) | 141.1 (8.034) | 0.6574 |
| **Potassium [mmol/L; median (IQ)]** | 4.205 (3.845-4.44) | 3.82 (3.673-4.17) | 4.365 (4.12-5.075) | 0.0022 (**) |
| **Chloride [mmol/L; mean (SD)]** | 100.9 (5.409) | 101.7 (3.851) | 100.4 (6.258) | 0.5591 |
| **Ferritin [mg/mL; median (IQ)]** | 758.3 (524.4-1.527) | 1.154 (629.2-1.513) | 594.2 (396.7-2.000) | 0.5952 |
| **Lactic dehydrogenase [U/L; mean (SD)]** | 501 (210.2) | 457.8 (171.9) | 525.6 (231.6) | 0.4795 |
| **C-Reactive Protein [mg/L; mean (SD)]** | 119.4 (85.17) | 82.34 (59.56) | 146 (92.54) | 0.0701 |
| **D-dimer [mg/L; median (IQ)]** | 1.715 (0.59-6.183) | 1.38 (0.525-2.885) | 2.05 (0.56-17.3) | 0.4912 |
| **Total leukocytes [cells/mm$^3$; mean (SD)]** | 13.776 (6.119) | 9.650 (2.932) | 16.355 (6.230) | 0.0041 (**) |
| **% Lymphocytes [median (IQ)]** | 9.050 (5.75-16.65) | 14.85 (10.3-22.5) | 6.95 (4.7-10.5) | 0.0048 (**) |

| | | | | |
|---|---|---|---|---|
| **Total Lymphocytes [cells/mm$^3$; median (IQ)]** | 1222 (751.6-1988) | 1419 (860.6-2525) | 1195 (658.4-1860) | 0.2199 |
| **% Neutrophils [median (IQ)]** | 81.7 (70.18-87.03) | 73.85 (66.55-82.23) | 86.1 (81.03-89.88) | 0.0139 (*) |
| **Neutrophil/Lymphocyte ratio [mean (SD)]** | 10.09 (7.097) | 5.277 (3.194) | 13.1 (7.26) | 0.0038 (**) |
| **Platelet count [cells/mm$^3$; mean (SD)]** | 241.323 (115.764) | 284.800 (120.550) | 214.150 (107.556) | 0.1327 |
| **Laboratory Findings - Blood Gas** | | | | |
| **pH [median (IQ)]** | 7.405 (7.371-7.444) | 7.435 (7.412-7.444) | 7.375 (7.352-7.439) | 0.0223 (*) |
| **$O_2$ Pressure [mmHg; median (IQ)]** | 68.7 (58.33-86.4) | 58.15 (50.95-65.15) | 80.1 (66.03-89.85) | 0.0028 (**) |
| **$CO_2$ Pressure [mmHg; mean (SD)]** | 44.28 (13.58) | 38.21 (4.91) | 48.08 (15.9) | 0.0704 |
| **$HCO_3$ [mmol/L; mean (SD)]** | 26.63 (4.91) | 24.84 (2.905) | 27.75 (5.625) | 0.1448 |
| **$CO_2$ Tension [mmol/L; median (IQ)]** | 22.1 (20.4-27.28) | 20.3 (19.6-24.15) | 23.25 (21.83-29.6) | 0.0309 (*) |
| **Excess Bases [mmol/L; mean (SD)]** | 1.4 (4.529) | 0.78 (2.408) | 1.788 (5.504) | 0.5915 |
| **$O_2$ Saturation [%; mean (SD)]** | 93.06 (4.756) | 90.25 (5.04) | 94.94 (3.609) | 0.0123 (*) |
| **Antivirals** | | | | |
| **Oseltamivir [n (%)]** | 19 (73.08) | 10 (100) | 9 (56.25) | 0.0227 (*) |
| **Antibiotics** | | | | |
| **Azithromycin [n (%)]** | 19 (73.08) | 8 (80) | 11 (68.75) | 0.668 |
| **Ceftriaxone [n (%)]** | 7 (26.92) | 3 (30) | 4 (25) | >0.9999 |
| **Clavulanate [n (%)]** | 11 (42.31) | 7 (70) | 4 (25) | 0.0426 (*) |
| **Amoxicillin [n (%)]** | 11 (42.31) | 7 (70) | 4 (25) | 0.0426 (*) |
| **Gentamicin [n (%)]** | 1 (3.85) | - | 1 (6.25) | >0.9999 |
| **Tazocin [n (%)]** | 1 (3.85) | - | 1 (6.25) | >0.9999 |
| **Vancomycin [n (%)]** | 1 (3.85) | - | 1 (6.25) | >0.9999 |
| **Polymyxin [n (%)]** | 1 (3.85) | - | 1 (6.25) | >0.9999 |
| **Anthelmintics** | | | | |
| **Ivermectin [n (%)]** | 4 (15.38) | 3 (30) | 1 (6.25) | 0.2642 |
| **Bacterial/Fungal Co-infections** | | | | |
| **Co-infection [n (%)]** | 14 (53.85) | 2 (20) | 12 (75) | 0.0138 (*) |

| | | | | |
|---|---|---|---|---|
| ***Candida* sp. [n (%)]** | 6 (23.08) | 1 (10) | 5 (31.25) | 0.3524 |
| ***Acinetobacter* sp. [n (%)]** | 5 (19.23) | 1 (10) | 4 (25) | 0.6169 |
| ***Enterococcus* sp. [n (%)]** | 5 (19.23) | - | 5 (31.25) | 0.1213 |
| ***Staphylococcus* sp. Coagulase- [n (%)]** | 5 (19.23) | 1 (10) | 4 (25) | 0.6169 |
| ***Staphyloccocus aureus* [n (%)]** | 3 (11.54) | - | 3 (18.75) | 0.2615 |
| ***Staphyloccocus aureus* MRSA [n (%)]** | 1 (3.85) | - | 1 (6.25) | >0.9999 |
| ***Pseudomonas aeruginosa* [n (%)]** | 4 (15.38) | - | 4 (25) | 0.1358 |
| ***Klebsiella pneumoniae* [n (%)]** | 5 (19.23) | - | 5 (31.25) | 0.1213 |
| ***Klebsiella ozanae* [n (%)]** | 1 (3.85) | - | 1 (6.25) | >0.9999 |
| ***Klebsiella oxytoca* [n (%)]** | 1 (3.85) | - | 1 (6.25) | >0.9999 |
| ***Serratia* sp. [n (%)]** | 2 (7.69) | - | 2 (12.5) | 0.5077 |
| **VRE [n (%)]** | 1 (3.85) | - | 1 (6.25) | >0.9999 |
| **KPC [n (%)]** | 1 (3.85) | - | 1 (6.25) | >0.9999 |
| ***Escherichia coli* [n (%)]** | 1 (3.85) | 1 (10) | - | 0.3846 |
| ***Proteus penneri* [n (%)]** | 1 (3.85) | - | 1 (6.25) | >0.9999 |
| ***Enterobacter* sp. [n (%)]** | 1 (3.85) | - | 1 (6.25) | >0.9999 |
| **Complications** | | | | |
| **Patients on mechanical ventilation at the time of blood collection [n (%)]** | 13 (50) | - | 13 (81.25) | 0.0001 (***) |
| **Severe Acute Respiratory Syndrome [n (%)]** | 25 (96.15) | 10 (100) | 15 (93.75) | >0.9999 |
| **Toxic shock syndrome [n (%)]** | 5 (19.23) | 1 (10) | 4 (25) | 0.6169 |
| **Renal Failure [n (%)]** | 4 (15.38) | - | 4 (25) | 0.1358 |
| **Pneumonia [n (%)]** | 5 (19.23) | 2 (20) | 3 (18.75) | >0.9999 |
| **Sepsis [n (%)]** | 3 (11.54) | - | 3 (18.75) | 0.2615 |
| **Respiratory Failure [n (%)]** | 3 (11.54) | - | 3 (18.75) | 0.2615 |
| **Cardiorespiratory Arrest [n (%)]** | 3 (11.54) | - | 3 (18.75) | 0.2615 |
| **Pulmonary embolism [n (%)]** | 1 (3.85) | - | 1 (6.25) | >0.9999 |
| **Hydroelectrolytic disorder [n (%)]** | 1 (3.85) | - | 1 (6.25) | >0.9999 |
| **Bradycardia [n (%)]** | 1 (3.85) | - | 1 (6.25) | >0.9999 |
| **Lymphocytopenia [n (%)]** | 1 (3.85) | - | 1 (6.25) | >0.9999 |
| **Outcome** | | | | |

| | | | | |
|---|---|---|---|---|
| **Hospital discharge [n (%)]** | 13 (50) | 9 (90) | 4 (25) | 0.0036(**) |
| **Death [n (%)]** | 13 (50) | 1 (10) | 12 (75) | |
| **Time between recruitment and patient outcome [days; median (IQ)]** | 8 (3.75-12) | 3 (1-6.25) | 10 (6.25-20.75) | 0.0013 (**) |
| **Total Length of Hospitalization [days; mean (SD)]** | 23.42 (12.15) | 16.2 (6.07) | 27.94 (12.93) | 0.0132 (*) |

**Table. S1. Patients' data.** Cardiovascular Diseases: Systemic Arterial Hypertension, Stroke, Acute Myocardial Infarction, Heart Failure, Aneurysm, Cardiopathic Patient. Pulmonary Diseases: Chronic Obstructive Pulmonary Disease, Bronchitis, Asthma. Neurological Conditions: Epilepsy/ Structural Epilepsy, Neurogenic Bladder, Diabetic Neuropathy, Alzheimer's Disease, Traumatic Brain Trauma, Paraparesis. Other Conditions: Hemochromatosis, Hypothyroidism, Motor Sequelae and Diabetic Retinopathy. Kidney Diseases: Chronic Kidney Disease. For parametric data, an unpaired Student's T test was used to determine statistical significance, for nonparametric data, Mann-Whitney test was used, both two-tailed. *P* values < 0.05 were considered statistically significant. * *P* < 0.05; ** *P* < 0.01; *P* < 0.001; *P* < 0.0001. The comparison between qualitative data was made with two-sided Fisher's exact test. The markers D-dimer, Ferritin and C-Reactive Protein presented values described as >32.5, >2000 and >320, respectively. These were considered as 32.5; 2000 and 32.5, respectively. One patient was excluded from glucose analysis and one from oxygen saturation analysis, as both were considered outliers (ROUT test (Q=1%)). Abbreviations: SD = Standard Deviation; IQ = Interquartile Range; MOD = moderate COVID-19; SEV = severe COVID-19; VRE = Vancomycin-Resistant enterococo; KPC = *Klebsiella pneumoniae carbapenemase*; MRSA = Methicillin-resistant *Staphylococcus aureus*.

| mRNA | Acession number | Sequence (5'->3') | |
|---|---|---|---|
| *RNA18SN1* | NR_145820.1 | Forward primer | CTCAACACGGGAAACCTCAC |
| | | Reverse primer | CGCTCCACCAACTAAGAACG |
| *CCL1* | NM_002981.2 | Forward primer | TGCAGATCATCACCACAGCC |
| | | Reverse primer | GTCCACATCTTCCGGCCA |
| *CCL2* | NM_002982.4 | Forward primer | CTCTGCCGCCCTTCTGTG |
| | | Reverse primer | TGCATCTGGCTGAGCGAG |
| *CCL3* | NM_002983.3 | Forward primer | AGCTGACTACTTTGAGACGAGCAG |
| | | Reverse primer | CGGCTTCGCTTGGTTAGGA |
| *CXCL9* | NM_002416.3 | Forward primer | TGCAAGGAACCCCAGTAGTGA |
| | | Reverse primer | GGTGGATAGTCCCTTGGTTGG |
| *IL1B* | NM_000576.3 | Forward primer | CAGAAGTACCTGAGCTCGCC |
| | | Reverse primer | CCTGGAAGGAGCACTTCATCT |
| *IL6* | NM_000600.5 | Forward primer | CTCCTTCTCCACAAGCGCC |
| | | Reverse primer | GATGCCGTCGAGGATGTACC |
| *IL17A* | NM_002190.3 | Forward primer | TCCCACGAAATCCAGGATGC |
| | | Reverse primer | GTCCTCATTGCGGTGGAGAT |
| *TNF* | NM_000594.4 | Forward primer | CTCTCTCTAATCAGCCCTCTGG |
| | | Reverse primer | CTCAGCTTGAGGGTTTGCTACAAC |
| *HAVCR2* | NM_032782.5 | Forward primer | CTACTGCTGCCGGATCCAAA |
| | | Reverse primer | GTGTCTGTGTCTCTGCTGGG |
| *CD4* | NM_000616.5 | Forward primer | ACAAGGAGGCAAAGGTCTCG |
| | | Reverse primer | CCATGTGGGCAGAACCTTGA |
| *CD8A* | NM_001768.7 | Forward primer | GCTGGACTTCGCCTGTGATA |
| | | Reverse primer | ACACGTCTTCGGTTCCTGTG |

**Table. S2. COVID-19 hyperinflammation-related genes panel primers.** Primers designed with Primer-BLAST (NCBI).

| Primer name | Sequence (5'->3') | | Predicted CpGs in product |
|---|---|---|---|
| CCL3_BC | Forward primer | TGTAGAGAGTTATGGTGTAGAGGAGG | 21 |
| | Reverse primer | CACCAAAAACCCTAAATTATACAAC | |

**Table. S3. BSP primers.** Primer designed with MethPrimer 2.0 software using the bisulfite converted target sequence GRCh38/hg38 chr17:36,090,276-36,090,005.
